# Scupa: Single-cell unified polarization assessment of immune cells using the single-cell foundation model

**DOI:** 10.1101/2024.08.15.608093

**Authors:** Wendao Liu, Zhongming Zhao

## Abstract

Immune cells undergo cytokine-driven polarization in respond to diverse stimuli. This process significantly modulates their transcriptional profiles and functional states. Although single-cell RNA sequencing (scRNA-seq) has advanced our understanding of immune responses across various diseases or conditions, currently there lacks a method to systematically examine cytokine effects and immune cell polarization. To address this gap, we developed **S**ingle-**c**ell **u**nified **p**olarization **a**ssessment (Scupa), the first computational method for comprehensive immune cell polarization analysis. Scupa is trained on data from the Immune Dictionary, which characterizes 66 cytokine-driven polarization states across 14 immune cell types. By leveraging the cell embeddings from the Universal Cell Embeddings model, Scupa effectively identifies polarized cells in new datasets generated from different species and experimental conditions. Applications of Scupa in independent datasets demonstrated its accuracy in classifying polarized cells and further revealed distinct polarization profiles in tumor-infiltrating myeloid cells across cancers. Scupa complements conventional single-cell data analysis by providing new insights into immune cell polarization, and it holds promise for assessing molecular effects or identifying therapeutic targets in cytokine-based therapies.

## Introduction

Immune cells detect and respond to a variety of stimuli, ensuring the body can effectively combat infections and other environmental or biological threats. Cytokines as crucial signaling molecules that facilitate communication between immune cells. These cytokines can induce significant changes in the transcriptional profiles and functional states of immune cells, a phenomenon known as immune cell polarization^1,2^. Through polarization, immune cells adapt their responses to better address specific challenges, enhancing the overall effectiveness of the immune system.

Recently, single-cell RNA sequencing (scRNA-seq) has been widely applied in immunological studies to investigate the responses of immune cells under various conditions. The production and response to cytokines, which play critical regulatory roles, have been extensively studied in numerous diseases, including COVID-19^3,4^, rheumatoid arthritis^5^, and cancers^6,7^. Importantly, a recent study systematically characterized the responses of 14 immune cell types to each of 86 cytokines and summarized the results as the Immune Dictionary^8^. A total of 66 cytokine-driven cell polarization states were identified in that study, serving as a valuable reference for assessing immune cell polarization in other scRNA-seq studies.

Currently, the investigation of immune cell polarization in scRNA-seq data is limited and there lacks consistent standards across studies. Most studies identified polarized cells based on the expression of certain signature genes from previous findings. However, this empirical approach suffers from technical noises, such as dropout effect and batch effect^9,10^, as well as biological variations including tissue or disease variations. For example, many studies analyzed macrophage polarization using M1 and M2 signature genes, but there was no consensus in the signature gene lists and the expression of these genes varied by conditions^11-13^. To address this challenge, recent advances in single-cell foundation models offer unprecedented potential. These models were trained on vast amounts of scRNA-seq data with millions of cells across all the representative human organs and learned cell representations within a unified biological latent space^14-16^. As they have been successfully demonstrated in multiple downstream tasks like cell type annotation and batch correction, we hypothesized that single-cell foundation models could also effectively represent immune cell polarization.

To facilitate the analysis of immune cell polarization in scRNA-seq data, we developed the method Scupa for **S**ingle-**c**ell **u**nified **p**olarization **a**ssessment. After being trained on scRNA-seq data from the Immune Dictionary, Scupa learns the representations of immune cell polarization within the latent space of Universal Cell Embeddings (UCEs)^16^. As the first computational method for systematic immune cell polarization analysis, Scupa enables the assessment of individual cell polarization across various predefined cytokine-driven cell polarization states in any scRNA-seq dataset.

## Results

### Overview of Scupa framework and immune cell polarization states

Immune cells undergo transcriptional and phenotypical changes in cytokine-driven polarization. Scupa uses the immune cell polarization states in the Immune Dictionary as the reference, and trains machine learning models to distinguish between polarized cells from cytokine-treated samples and unpolarized cells from phosphate buffered saline (PBS)-treated control samples. The specific polarized states of a cell type are usually driven by the specific cytokines, with each state exhibiting a unique transcriptional profile. Instead of relying on gene expression features, which can vary across species, tissues, and conditions, Scupa utilizes cell embeddings from the single-cell foundation model UCE for assessing immune cell polarization^16^. We chose UCE over other single-cell foundation models because of its advantages in multiple-species compatibility, and ‘zero-shot’ capability without the need for additional fine-tuning. We further reduced the dimension of cell embeddings using principal component analysis (PCA), and trained support vector machine (SVM) models to classify polarized cells and unpolarized cells based on principal components (PCs). The models learned the representation of each polarized state in the latent space of UCEs, allowing for transferability to other datasets (Fig. 1a). We found that SVM outperformed several other machine learning models in this task (Supplementary Table 1).

**Fig. 1.**
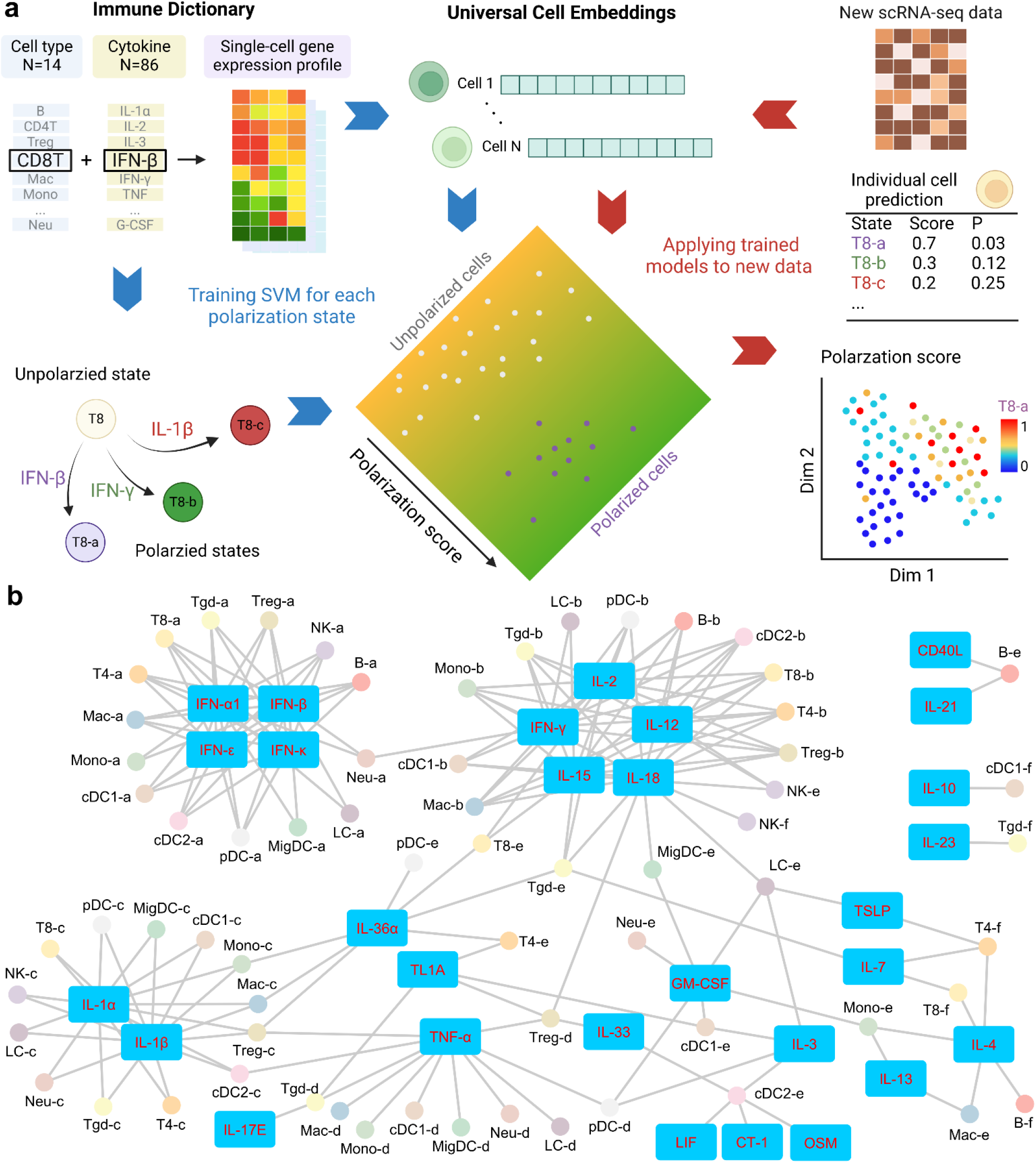
The framework of Scupa and cytokine-driven cell polarization states. a. Scupa uses the Immune Dictionary as the reference and training set for measuring immune cell polarization. Major immune cell polarization states were defined in the Immune Dictionary according to driving cytokines and top marker genes. Universal Cell Embeddings (UCEs) were generated for the Immune Dictionary and any new scRNA-seq dataset for Scupa. Scupa trained support vector machine (SVM) models for various cell polarization states using the Immune Dictionary, and then it applies the trained models to predict polarization in a new dataset. It outputs the polarization score and p-value for each individual cell. Created with BioRender.com. b. A network showing the driving cytokines and immune cell polarization states.

For any new scRNA-seq dataset containing immune cells presented in the Immune Dictionary, Scupa examines whether the cells have the similar transcriptional changes as reference polarized cells, thereby inferring their polarization states and received cytokines. The polarization of each cell is assessed based on its cell embeddings from UCE. The trained SVM models predict the polarization scores of each cell using the learned representation in the unified latent space of UCEs. According to the cell type, Scupa assigns a score to each individual cell for every polarization state, ranging from 0 (unpolarized) to 1 (fully polarized). Additionally, Scupa outputs a p-value by comparing the polarization score of a cell with the null distribution of polarization scores from the reference unpolarized cells, facilitating the identification of significantly polarized cells. Scupa is designed for integration into the widely used Seurat pipeline for comprehensive single-cell data analyses^17^, enabling the output scores and p-values to be readily visualized in multiple formats (Fig. 1a).

Scupa supports the analysis of 66 polarization states in 14 immune cell types from the Immune Dictionary (Fig. 1b, Supplementary Fig. 1). Among these states, the ‘a states’ of all cell types represent those driven by some type-I interferons (IFN-α1, IFN-β, IFN-ε, IFN-κ). The ‘b states’ represent those driven by IFN-γ and interleukins inducing IFN-γ expression (IL-2, IL-12, IL-15, IL-18). The ‘c states’ are for those driven by two proinflammatory cytokines IL-1α and IL-1β, and the ‘d states’ are mainly driven by TNF-α. In contrast, the driving cytokines of the ‘e states’ and ‘f states’ in various cell types tend to vary. Identification of polarized cells using Scupa suggests the potential presence of one or more driving cytokines in the tissue, thereby facilitating the understanding of immune cell environment, communication and response in scRNA-seq data.

### Scupa learns the representation of cell polarization in various immune cell types

Cytokines are key regulators of intracellular signaling and gene expression, acting as messengers that mediate and modulate immune responses. They activate signaling cascades that lead to the phosphorylation and activation of various transcription factors, which then translocate to the nucleus to modulate the transcription of genes. Consequently, the cytokine-driven immune cell polarization states are characterized by unique transcriptional profiles^8^. UCE can effectively capture these transcriptional changes and represent them as variations in cell embeddings. For example, we found that the CD8^+^ T cells treated with driving cytokines of each polarization state tended to have cell embeddings shifting away from the unpolarized cells, as visualized by uniform manifold approximation and projection (UMAP, Fig. 2a). By filtering cells based on cosine similarity and the expression of top marker genes, we obtained fully polarized cells in each state, which displayed distinctly different cell embedding distributions from unpolarized cells (Fig. 2b).

**Fig. 2.**
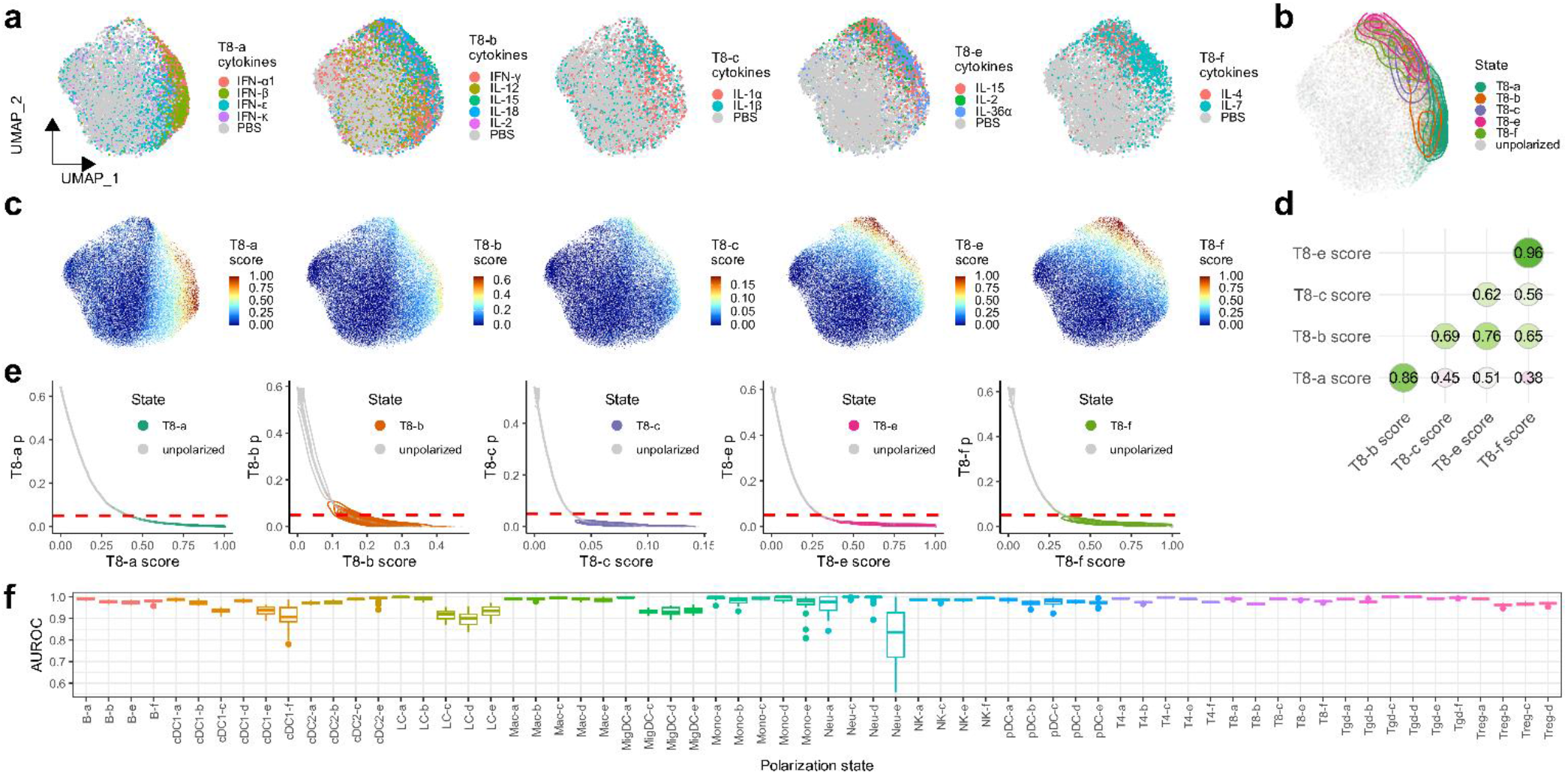
Scupa learns the representation of polarization states in CD8^+^ T cells and other cells. a. Uniform manifold approximation and projection (UMAP) plots showing CD8^+^ T cells from the samples treated with driving cytokines of each polarization state or from the control samples treated with PBS. UMAPs are derived from UCE cell embeddings rather than gene expression. b. UMAP plot showing the distribution of fully polarized cells of each polarization state after filtering. c. UMAP plots showing polarization scores of each polarization state from Scupa prediction. d. The Spearman correlation coefficients between polarization scores from each two polarization states. e. The polarization scores and p-values of fully polarized cells of each polarization state and unpolarized cells. The red dashed line indicates p-value=0.05. f. Box plots showing the testing AUROC values in 20 repeats across all polarization states.

For each polarization state, we trained an SVM model to classify polarized cells and unpolarized cells, and quantified the polarization using a score derived from the trained models. With this approach, the gradients of polarization scores in the unified cell embedding space represent the directions of cell polarization (Fig. 2c). Among all CD8^+^ T cells, we found that the polarization scores of some states are highly correlated (Fig. 2d). For example, the Spearman correlation coefficient between T8-e and T8-f state scores is 0.96. This indicates a high similarity in transcriptional changes between these two states, which aligns with the findings in the original study^8^. We then calculated the p-values for each polarization state by comparing the polarization scores with the distribution of scores from unpolarized cells. This metrics is useful because the commonly applied p-value threshold of 0.05 can serve as a good criterion for identifying fully polarized cells across most polarization states (Fig. 2e). Importantly, in addition to CD8^+^ T cells, we observed similar patterns of polarization scores and p-values across all other 13 cell types and 61 polarization states. This demonstrated that Scupa could effectively learn the representation of all polarization states in the unified cell embedding space (Supplementary Figs. 2-14).

Next, we evaluated the performance of Scupa across all polarization states. Using a random 70% of the data for training and the remaining 30% for testing, we repeated this process 20 times and calculated the area under the receiver operating characteristic curve (AUROC) for testing set. The median AUROC values were above 0.95 in 56 out of the 66 polarization states (Fig. 2f). For polarization states with lower performance, such as Neu-e (AUROC = 0.836), the primary factor was the insufficient number of cells leading to poor model fitting (Supplementary Fig. 15). Overall, Scupa achieves superior performance in learning the representations of polarization states in scRNA-seq data.

### Scupa characterizes cytokine-driven immune cell polarization in both *in vitro* and *in vivo* datasets

To evaluate Scupa’s performance in independent datasets, we collected two scRNA-seq datasets generated from cytokine-treated samples. The first dataset comprises human peripheral blood mononuclear cell (PBMC) samples treated with IFN-β or left untreated *in vitro*^18^. IFN-β treatment dramatically induces the transcription of interferon-stimulated genes, causing the cell embeddings of treated cells to diverge significantly from those of untreated cells across all immune cell types (Fig. 3a). We applied Scupa to analyze the immune cell polarization in this dataset. Considering that IFN-β is the driving cytokine of -a states in all immune cell types (Fig. 1b), we examined the polarization scores of these states. We found that all cells from the IFN-β-treated sample had polarization scores close to 1, while those cells from the untreated sample had scores close to 0. This clearly indicates a tremendous difference (Fig. 3b). Additionally, nearly all cells from the IFN-β-treated sample had polarization p-values below 0.05, whereas those cells from the untreated sample had p-values above 0.05 (Fig. 3c, Supplementary Fig. 16). The ROC curves further confirmed the near-perfect performance of using polarization scores to classify cells from treated and untreated samples, with AUROCs above 0.99 across all cell types (Fig. 3d). Importantly, although Scupa was trained on mouse scRNA-seq data, it demonstrated excellent performance on this human scRNA-seq data without any adaptation, suggesting that Scupa learned unified representations of immune cell polarization across species.

**Fig. 3.**
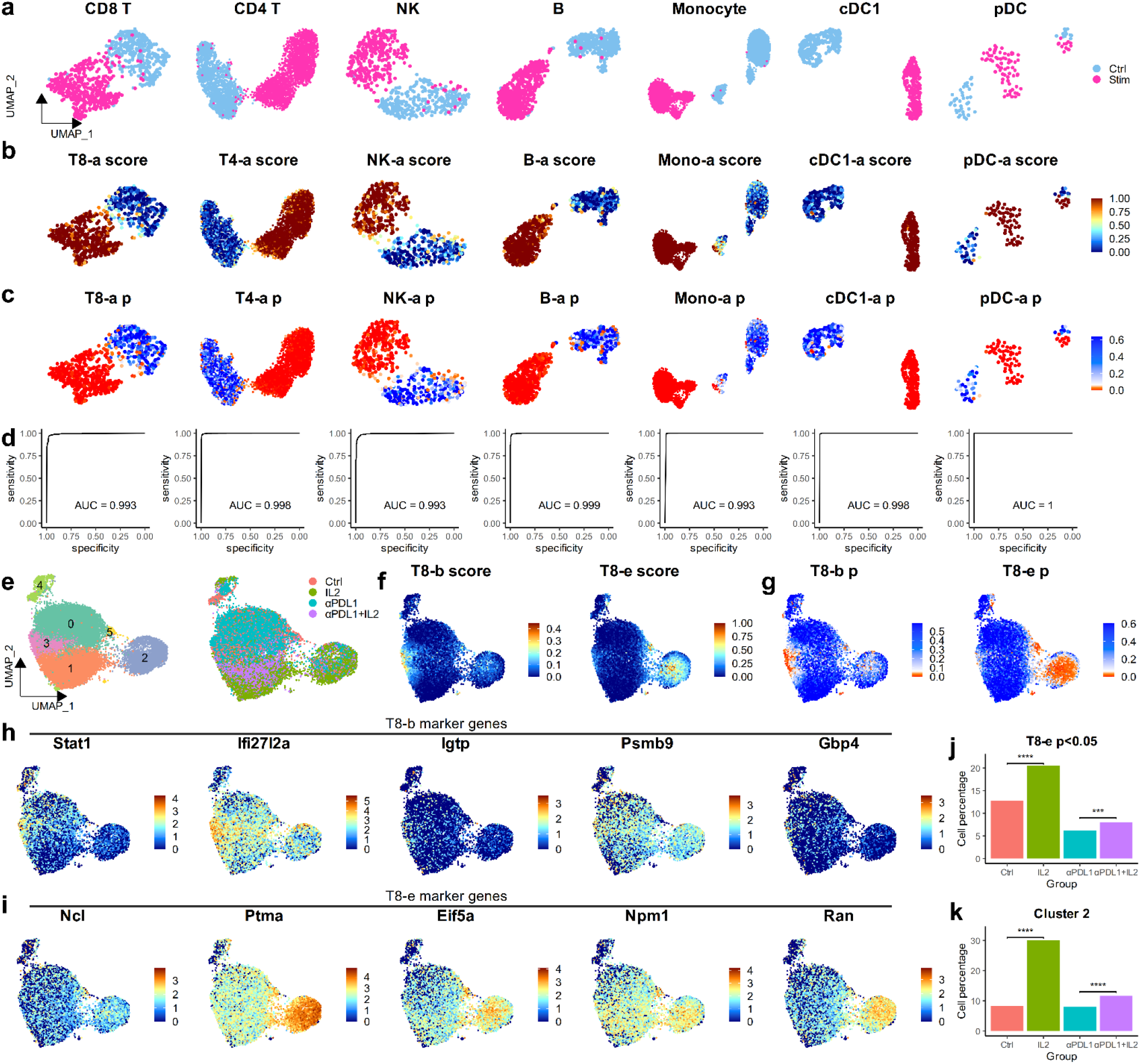
Validation of Scupa in *in vitro* and *in vivo* datasets. a. UMAP plots showing the immune cells from the control and IFN-β-stimulated samples. UMAPs are derived from UCE cell embeddings rather than gene expression. Ctrl: untreated control sample. Stim: interferon-stimulated sample. b. UMAP plots showing the polarization scores in different immune cell types. The cells from IFN-β-stimulated samples generally had much higher scores. c. UMAP plots showing the polarization p-values in different immune cell types. The cells from IFN-β-stimulated samples generally had much lower p-values. d. ROC curves showing the superior performance of polarization scores for classifying immune cells from the control and IFN-β-stimulated samples. e. UMAP plots showing CD8+ T cells in different clusters and treatment groups. f. UMAP plots showing polarization scores of two IL-2-driving states, T8-b and T8-e. g. UMAP plots showing polarization p-values of two IL-2-driving states. h. The expression level of five T8-b marker genes. i. The expression level of five T8-e marker genes. j. The percentage of cells with T8-e polarization p-value<0.05 among different treatment groups. k. The percentage of cells in cluster 2 among different treatment groups. ***: p<0.001. ****: p<0.0001.

In the second study, the mice chronically infected with lymphocytic choriomeningitis virus (LCMV) were treated with IL-2, anti-PD-L1, or a combination therapy^19^. The virus-specific CD8^+^ T cells from the mouse spleen were sorted for scRNA-seq. We clustered cells from three treatment groups and control group using UCE cell embeddings (Fig. 3e), and applied Scupa to analyze the CD8^+^ T cell polarization in this dataset. As IL-2 is the driving cytokine for two polarization states, T8-b and T8-e, we examined the polarization scores and p-values of these two states (Fig. 3f, g). A small number of cells were polarized to T8-b state and enriched in cluster 3, while more cells were polarized to T8-e state and enriched in cluster 2. To verify the two polarization states with shared driving cytokines, we examined the expression of polarization state marker genes identified in the Immune Dictionary^8^. We found that the expression of some marker genes correlated with polarization scores, such as *Stat1* with T8-b scores and *Ptma* with T8-e scores. This result suggested that Scupa could distinguish these two polarization states. On the other hand, we observed the ubiquitous high or low expression of some marker genes (e.g., *Igtp, Gbp4, Ncl*, and *Npm1*) in all CD8^+^ T cells, highlighted the variability of gene expression in different datasets (Fig. 3h, i). Therefore, the feature of independence on marker genes in Scupa makes it more applicable to various scRNA-seq data from diverse sources.

Since IL-2 was administrated *in vivo*, it was likely that only a subset of spleen T cells were polarized by IL-2, while other T cells were polarized by different cytokines or remained unpolarized. To analyze the effect of IL-2 treatment, we compared the percentage of cells with significant T8-e polarization (polarization p-value<0.05) among groups. This percentage significantly increased in the IL-2 treatment group when compared to the control group (p<0.0001), and increased in the combination therapy treatment group when compared to the anti-PD-L1 treatment group (p<0.001, Fig. 3j). We also compared the percentage of cells in cluster 2, which was highly enriched with the cells polarized to T8-e state. Similar increase was observed with IL-2 treatment (Fig. 3j), confirming the CD8+ T cell polarization to IL-2 driving polarization states.

As above, Scupa was validated using scRNA-seq datasets generated from both *in vitro* and *in vivo* samples. It superbly classified stimulated and unstimulated cells from *in vitro* samples, and also revealed the increases of cell polarized to a cytokine-driving polarization state in *in vivo* samples.

### Scupa reveals polarization states and proinflammatory responses of myeloid cells across cancer types

Myeloid cells play a crucial role in the tumor microenvironment, influencing cancer progression and response to therapy. These cells, which include macrophages, monocytes, dendritic cells (DCs), and neutrophils, can exhibit either pro-tumor or anti-tumor functions depending on their polarization state and the cytokine milieu^20,21^. Understanding the dual roles of myeloid cells in cancers is essential for developing therapeutic strategies that modulate their function to inhibit tumor progression and improve patient outcomes. In order to systematically analyze the myeloid cell polarization in multiple cancer types, we applied Scupa to a pan-cancer single-cell atlas of tumor infiltrating myeloid cell dataset^11^.

We first revisited the macrophage polarization using Scupa as it has been extensively investigated in cancer research^22^. We calculated the polarization scores of five polarization states for all macrophage clusters and compared these scores across seven cancer types. Among the five states, Mac-b, Mac-c, and Mac-d are all M1-like states driven by proinflammatory cytokines, while Mac-e is a M2-like state driven by cytokines that induce M2 polarization (Fig. 1b)^2^. Several macrophage clusters displayed consistent polarization profiles across cancer types. *C1QC*^+^ and *LYVE1*^+^ macrophages generally had low polarization scores of all polarization states, indicating that they were mainly unpolarized. *ISG15*^+^ macrophages generally had high Mac-a, Mac-b, Mac-c, and Mac-d polarization scores, suggesting that they were primarily polarized by type-I interferons and proinflammatory cytokines. *NLRP3*^+^ and *INHBA*^+^ macrophages also had high Mac-b, Mac-c, and Mac-d scores. This might indicate polarization by proinflammatory cytokines. In particular, *SPP1*^+^ macrophages showed significant variation across these cancer types. They had highest Mac-e polarization scores in thyroid carcinoma (THCA), but had much lower scores in pancreatic adenocarcinoma (PAAD) or uterine corpus endometrial carcinoma (UCEC). When comparing macrophage polarization in different cancer types, we found that PAAD and kidney cancer had the lowest overall polarization scores in all polarization states. This is potentially due to lower cytokine production in these tumors (Fig. 4a).

**Fig. 4.**
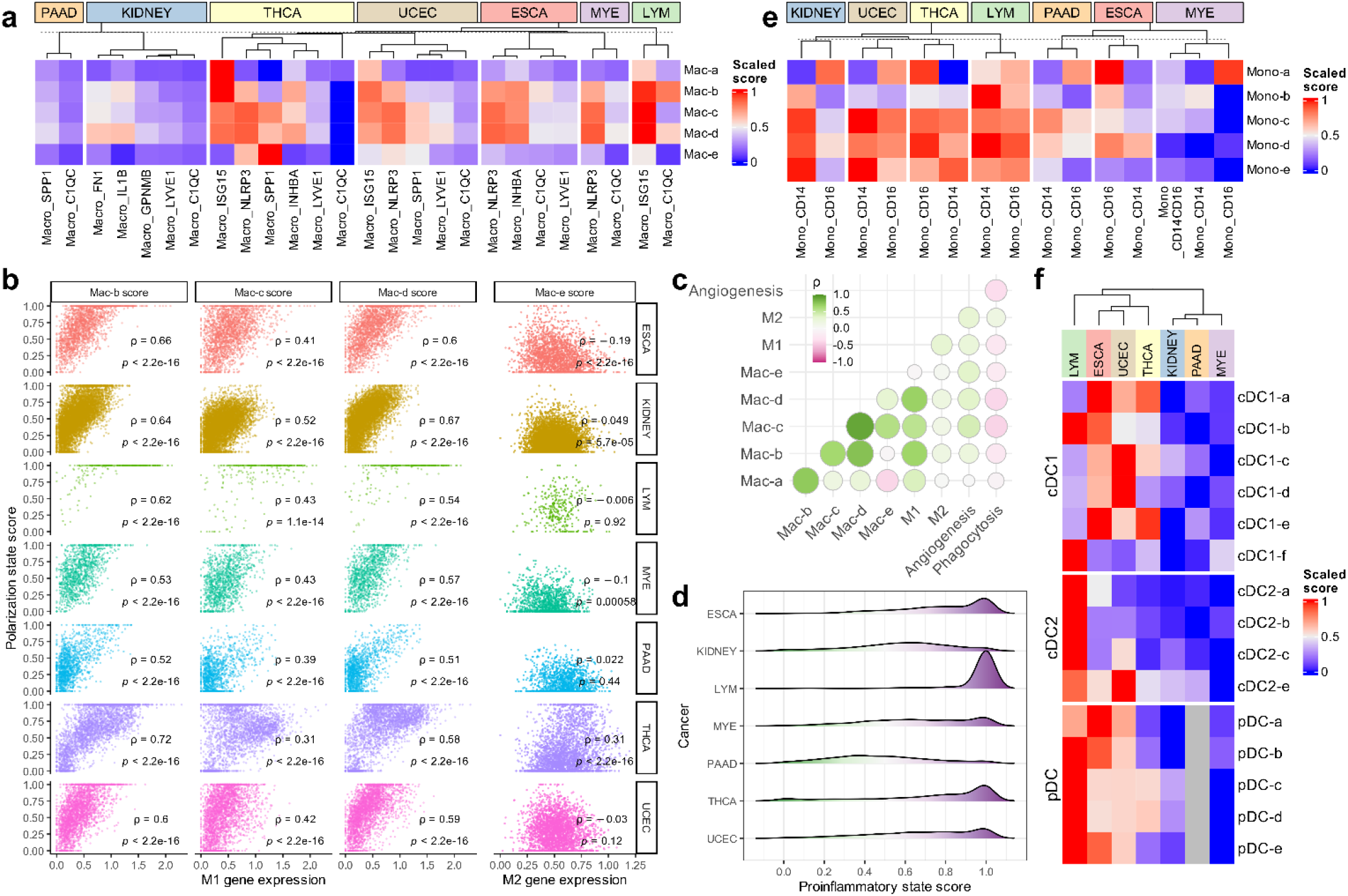
Comparison of the polarization states of infiltrating myeloid cells across seven cancer types by Scupa. a. The mean polarization scores in different macrophage subpopulations from different cancer types. Polarization scores are scaled across cancer types. b. The polarization scores of macrophage states Mac-b, Mac-c, and Mac-d were positively correlated with M1 signature gene expression. In contrast, the polarization scores of macrophage state Mac-e were not positively correlated with M2 signature gene expression. c. The correlation between macrophage state polarization scores, M1 signature gene expression, M2 signature gene expression, angiogenesis signature gene expression, and phagocytosis signature gene expression in all macrophages. d. LYM macrophages displays overall highest proinflammatory state scores. e. The mean polarization scores in different monocyte subpopulations from seven cancer types. f. The mean polarization scores in different DC subpopulations from seven cancer types. ESCA: esophageal carcinoma, KIDNEY: kidney cancer, LYM: lymphoma, MYE: myeloma, PAAD: pancreatic adenocarcinoma, THCA: thyroid carcinoma, UCEC: uterine corpus endometrial carcinoma.

Next, we analyzed the relationship between polarization scores of macrophage polarization states and the expression of M1 and M2 signature genes^11^. The mean expression of M1 signature genes was positively correlated with Mac-b, Mac-c, and Mac-d polarization scores across all cancer types. In contrast, there was no consistent correlation between the mean expression of M2 signature genes and Mac-e polarization scores (Fig. 4b). This lack of correlation was likely attributed to the observation of that the M2 signature genes did not overlap with the Mac-e marker genes from the Immune Dictionary, thus resulting in strong variation in different studies. Additionally, we included angiogenesis and phagocytosis signature genes in the correlation analysis^11^. The mean expression of angiogenesis signature genes was moderately positively correlated with Mac-b (Spearman’s correlation coefficient ρ=0.22), Mac-c (ρ=0.42), Mac-d (ρ=0.35), and Mac-e (ρ=0.35) polarization scores, highlighting macrophages’ simultaneous contributions to inflammation and angiogenesis^23^. Conversely, the mean expression of phagocytosis signature genes was negatively correlated with polarization scores of all polarization states, suggesting that phagocytotic macrophages were generally less polarized by cytokines (Fig. 4c). Considering the high correlation between Mac-b, Mac-c, and Mac-d polarization scores, and proinflammatory cytokines as driving cytokines for these states, we defined a macrophage proinflammatory state score as the maximum of these three scores in each single cell. This score summarizes macrophage proinflammatory polarization and its distribution indicates the proinflammatory activity in different cancers. We found that lymphoma (LYM) was characterized with the highest overall proinflammatory scores, followed by THCA and esophageal carcinoma (ESCA). PAAD and kidney cancer exhibited the lowest overall proinflammatory scores (Fig. 4d).

In addition to macrophages, we examined the polarization of monocytes and different DC populations. These cell populations displayed stronger variation in polarization across cancer types than macrophages (Fig. 4e, f). In LYM, *CD14*^+^ monocytes, *CD16*^+^ monocytes, cDC2, and pDCs had high polarization scores of all polarization states, supporting the strong effect and important role of multiple cytokines in the cancer^24^. In ESCA, UCEC, and THCA, monocytes, cDC1, and pDCs were also polarized to multiple states. Notably, the overall polarization of macrophages, monocytes, and DCs showed a consistent trend in all seven cancer types. All these myeloid cell populations were more polarized in LYM, THCA, UCEC, and THCA, but less polarized in PAAD, LYE, and kidney cancer. This trend likely reflects the variation in cytokine milieu in the tumor microenvironment of these cancer types (Fig. 4d, e, f). Taken together, these results indicated the distinct cytokine environments, myeloid cell polarization patterns, and proinflammatory responses in different cancers, demonstrating Scupa’s capability to complement the conventional scRNA-seq analysis with unique cell polarization analysis.

## Discussion

In this study, we introduce Scupa, the first computational method designed to assess immune cell polarization from scRNA-seq data. Scupa is design to complement to the conventional scRNA-seq analysis pipelines by providing additional perspectives in the cytokine environment and immune cell polarization. The method leverages the recently released Immune Dictionary, which systematically characterized the responses of 14 immune cell types to 86 cytokines and then identified 66 cytokine-driven polarization states. Unlike traditional approaches that rely on predefined signature genes, Scupa utilizes cell embeddings from the single-cell foundation model, Universal Cell Embeddings, to capture the nuanced transcriptional changes associated with different polarization states. Our results clearly indicated that Scupa could effectively classify polarized and unpolarized cells by training machine learning models on cell embeddings. The method was validated using independent datasets, including human PBMCs treated with IFN-β and CD8+ T cells from mice spleen treated with IL-2 and anti-PD-L1. Scupa accurately identified polarized cells and revealed the cytokine-driven polarization states within these datasets. With UCE’s multiple-species compatibility, even though our method was trained on the mouse data of Immune Dictionary, it could effectively be applied to human dataset without additional fine tuning. While this needs further evaluation, it suggested the robustness of our model across species.

Additionally, we apply Scupa to a pan-cancer single-cell atlas to investigate the polarization of myeloid cells across seven cancer types. The analysis reveals distinct polarization profiles and proinflammatory responses in macrophages, monocytes, and DC populations. Notably, LYM, THCA, UCEC exhibit higher polarization scores when compared to PAAD, LYE, and kidney cancer, reflecting distinct cytokine environments in different cancer types.

Macrophage polarization has been one of the top research interests in immunology due to its significant role in health and disease. Traditionally, macrophages have been categorized into two main polarization states. The first state is M1, or classically activated macrophages that are induced by proinflammatory cytokines like IFN-γ and TNF-α. These macrophages are associated with antimicrobial and tumoricidal activities. The second state is M2, or alternatively activated macrophages that are stimulated by cytokines such as IL-4 and IL-13. These macrophages are involved in tissue repair and immune regulation^2,22^. However, this dichotomous classification has been increasingly recognized as an oversimplification, with emerging evidence suggesting a spectrum of intermediate states influenced by a variety of cytokines and environmental cues^25^. In many scRNA-seq studies, the lack of consistent marker genes and standards further makes the polarization inference arbitrary. Scupa provides a signature gene-free approach for analyzing macrophage polarization to five cytokine-driving states. Using Scupa, we identified the *C1QC*^+^ or *LYVE1*^+^ unpolarized macrophage subpopulations and multiple macrophage subpopulations polarized to M1-like polarization states. Of note, Scupa analysis revealed *SPP1*^+^ macrophages polarized to Mac-e state, a M2-like polarization state, suggesting its pro-tumor roles. This result was further supported by previous findings that the worse clinical outcomes were associated with higher *SPP1* expression^11^.

Cytokine therapies have emerged as a powerful strategy for treating various diseases, leveraging the potent regulatory effects of cytokines to modulate immune responses and target disease processes. These therapies harness the ability of cytokines to influence cellular behavior, enhancing or suppressing immune responses as needed. For instance, cytokines such as interferons have been used to treat viral infections and certain cancer types^26,27^, while interleukins have shown promise in enhancing immune responses in cancer immunotherapy^19,28^. Scupa’s ability to systematically assess immune cell polarization based on scRNA-seq data offers a valuable tool for assessing the molecular effects of cytokine therapies (Fig. 3e-k). By evaluating how cytokine treatments impact the polarization and functional states of immune cells, Scupa can provide insights into the therapeutic mechanisms at a granular level. It also holds the potential to identify therapeutic targets for cytokine therapies, facilitating the development and optimization of cytokine-based treatments.

There are some limitations in this work. First, there lacks high-quality experimentally generated immune cell polarization states. Although we included all predefined polarization states from the Immune Dictionary, there may be additional, unidentified polarization states due to the constraints of experimental design. For example, some chemokines have been found to induce macrophage polarization^29,30^, but the chemokine family was not included in the Immune Dictionary. Second, there are noncytokine pathways of immune cell polarization, such as hypoxia and lactate for macrophage polarization^31,32^. Identification of new immune cell polarization states will require further investigations comparable to the Immune Dictionary, and necessary immunological expertise. Such data has not been available yet, but Scupa can be updated to include newly identified polarization states in future studies. Third, while Scupa has been demonstrated robust in identifying polarization and states in both mice and humans, more evaluation will be needed to enhance the models across species.

In conclusion, we introduced Scupa, the first method for comprehensive immune cell polarization assessment using scRNA-seq data. It is broadly appliable to the studies of various diseases involving immune cell populations and is particularly useful in contexts where cytokines play important roles in disease pathogenesis, progression, and treatment.

## Methods

### Generating cell embeddings using UCE and dimension reduction

We used single-cell foundation model, Universal Cell Embeddings (https://github.com/snap-stanford/UCE), to generate cell embeddings for all scRNA-seq datasets used in this study. The pretrained 4-layer model was employed with a batch size of 50. The UCE cell embeddings are 1,280-dimensional, representing cells in the unified latent space. However, this high dimensionality poses challenges for training machine learning models on most polarization states with a limited number of cells.

To address this issue, we performed PCA to reduce the dimensionality of UCE cell embeddings for each cell type. Principal components (PCs) are linear combinations of vector bases in the cell embedding space, with the top PCs representing directions with the largest variation for that cell type. By default, Scupa uses the first 20 PCs as features for machine learning training and prediction. Additionally, we generated two-dimensional UMAPs using the first 20 PCs for data visualization.

### Identifying fully polarized cells in the Immune Dictionary

According to both our analysis and the original study^8^, only a subset of immune cells were polarized after cytokine treatment in the *in vivo* experiments. This was likely due to variable cytokine concentration, receptor expression, and cellular status of different cells. The cell embeddings of some cells from cytokine-treated samples were closer to those of unpolarized cells from the PBS-treated samples than other cells from cytokine-treated samples, likely suggesting an unpolarized state or mildly polarization state (Fig. 2e, Supplementary Fig. 2-14). Therefore, we first identified fully polarized cells of each polarization state for training machine learning models.

We identified the fully polarized cells of each polarization state based on following three criteria. 1) The cell is from a sample treated with one of the driving cytokines. 2) The mean expression of top marker genes of the polarization in the cell is higher than that of most other cells. 3) The UCE cell embeddings of the cell are similar to those of the other cells from the samples treated with driving cytokines.

For criterion 2, we used a consistent threshold of the 90^th^ quantile, i.e., the mean expression of top marker genes in the cell was required to be higher than that in 90% cells of the same cell type. For criterion 3, we found that the UCE cell embeddings of all cells of the same cell type were highly correlated and, thus, not informative for identifying fully polarized cells. To overcome this issue, we calculated the ‘embedding shift’ as the vector difference between the cell embedding of each cell and the cell embedding of unpolarized cell center, which represented the cell embedding change from the unpolarized state. We then calculated the cosine similarity between the embedding shift of each cell with the rest cells from the samples treated with driving cytokines. Fully polarized cells had to secure a minimum mean cosine similarity, ranging from 0.08 to 0.2 among different cell types. Those cells that satisfied with all criteria were considered fully polarized cells and then were used for training machine learning models alongside unpolarized cells from PBS-treated samples.

### Training and testing machine learning models

When training machine learning models to classify unpolarized cells and polarized cells, we tested several models including: 1) logistic regression using the ‘glm’ function from R package ‘stats’, 2) SVM using the ‘svm’ function from R package ‘e1071’, 3) random forest using the ‘randomForest’ function from R package ‘randomForest’, and 4) semi-supervised learning approach. For each cell type, 70% cells were randomly selected for training and the remaining 30% for testing. During training, unpolarized cells were labeled with a polarization score of 0, while the fully polarized cells identified in the previous step were labeled with a polarization score of 1. For binary classification models, the predicted probability to the fully polarized state was used as the polarization score. For regression models, the output prediction was clamped to a range of 0 to 1 and used as the polarization score. We repeated the training and testing for 20 times and calculated the mean AUROC values for each machine learning model. SVM showed the best performance with the highest mean AUROC values across all polarization states. The final SVM models in Scupa were trained using the ‘svm’ function from R package ‘e1071’, with a linear kernel and eps-regression type^33^.

For the semi-supervised learning approach, we included the cells from the samples treated with cytokines other than the polarization state-driving cytokines as unlabeled data. We first trained supervised machine learning models (logistic regression, SVM, random forest) on labeled data: unpolarized cells and fully polarized cells from the previous step. The trained models were then used to classify unlabeled cells as either unpolarized or polarized. In the end, the final machine learning models were trained on the combined data from initial identification and following prediction. In our comparison of the testing results from supervised models with semi-supervised models, we found that the semi-supervised models generally had slight worse performance compared to the corresponding supervised models, despite that they improved the performance on some polarization states with small cell numbers (Supplementary Table 1). Therefore, we did not used the trained semi-supervised models for prediction in the Scupa package.

### Estimating statistical significance of polarization

When calculating the p-value of a cell for being polarized to a polarization state, we set the null hypothesis as H0: the observed cell is an unpolarized cell, and the alternative hypothesis as H1: the observed cell is a polarized cell. The polarization score serves as the test statistic. We used the polarization score distribution from unpolarized cells of a specific cell type in the Immune Dictionary as the null distribution. The p-value is calculated as the probability of obtaining a polarization score equal or greater than the observed polarization score from the null distribution. This hypothesis testing is a non-parametric, with no assumptions about the cell embedding distributions of the input dataset.

Scupa adjusts the p-values using the Benjamini-Hochberg procedure for the input dataset. We found that both unadjusted p-values and adjusted p-values accurately classified polarized cells in the samples from *in vitro* experiments (Supplementary Fig. 16). However, there were significantly fewer cells with adjusted p-values <0.05 in the samples from *in vivo* experiments. This discrepancy is likely due to varying extents of immune cell polarization across different studies, and possibly more dynamic natures in the living cells. Multiple experimental factors can influence the cytokine concentration in the tissue, thereby affecting the extent of immune cell polarization. Consequently, using a stringent adjusted p-value threshold for identifying fully polarized cells would result in few positive cells if cytokine concentration were low. Based on our analysis of four datasets included in this study, it is recommended to use unadjusted p-values<0.05 for identifying polarized cells in the samples from *in vivo* experiments.

### Cross-dataset batch effect correction

As a single-cell foundation model, UCE is robust to dataset and batch-specific artifacts, though cross-dataset batch effects may still persist between the Immune Dictionary and other datasets. To enhance Scupa’s transferability to diverse datasets, we provide a straightforward and effective approach for cross-dataset batch effect correction. UCE’s capability allows us to represent cross-dataset batch effects as the difference in unpolarized cell embeddings between two datasets. When there are untreated control and treated samples, cells from the control samples could be specified as unpolarized cells. Scupa first calculates the center of reference unpolarized cells’ UCE cell embeddings ***c***_*ref*_, and the center of input unpolarized cells’ UCE cell embeddings ***c***_*in*_. For a cell k with UCE cell embeddings (***emb***_*k*,*original*_) from the input dataset, its cell embeddings are adjusted to:

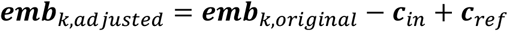

This adjustment allows the learned representations of immune cell polarization to be applied to the adjusted cell embeddings, bypassing complicated data integration processes and thus, preserving polarization information that might be lost with scRNA-seq data integration methods^34,35^. In the Scupa package, we implement this batch effect correction approach and provide an optional parameter for users to specify unpolarized cells in the input dataset, suitable for experimental designs with untreated healthy controls.

Regarding the two cytokine treatment datasets in Scupa evaluation, we used this batch effect correction approach when analyzing immune cell polarization. In the IFN-β treated human PBMC scRNA-seq dataset, the cells from the untreated sample were specified as unpolarized cells. Similarly, in the IL-2 treated mouse spleen scRNA-seq dataset, the cells from the untreated mouse were specified as unpolarized cells. We found only slight differences in the polarization analysis results when specifying unpolarized cells or not in both datasets. This evaluation indicated the robustness of UCE and Scupa to cross-dataset batch effect correction. For the pan-cancer myeloid cell dataset, we did not indicate unpolarized cells due to the absence of an untreated healthy control sample in the dataset.

### Statistical analysis

The calculation of p-values and adjusted p-values in Scupa is described in subsection “Deriving statistical significance of polarization” above. When comparing the cell proportions between two conditions in the IL-2 treated mouse spleen scRNA-seq dataset, we used two-sided Fisher exact test. All statistical analyses were performed in R (v 4.1.3).

## Data availability

All data used in the study are from published studies. The Immune Dictionary scRNA-seq dataset was downloaded from Single Cell Portal (https://singlecell.broadinstitute.org/single_cell/study/SCP2554/). The IFN-β treated human PBMC scRNA-seq dataset with cell type annotation was downloaded using SeuratData (https://github.com/satijalab/seurat-data). The rest datasets were downloaded from Gene Expression Omnibus (GEO) database with following accession numbers: IL-2 treated mouse spleen scRNA-seq dataset (GEO206732) and pan-cancer infiltrating myeloid cell scRNA-seq data (GSE154763). The processed datasets with generated UCE cell embeddings are available at https://zenodo.org/doi/10.5281/zenodo.13312248.

## Code availability

Scupa is available as a R package. The source code and vignettes of Scupa are freely available at https://github.com/bsml320/Scupa. The code for generating the results is available at https://zenodo.org/doi/10.5281/zenodo.13312248.

## Acknowledgements

W.L. is a CPRIT Predoctoral Fellow in the Biomedical Informatics, Genomics and Translational Cancer Research Training Program (BIG-TCR) funded by Cancer Prevention & Research Institute of Texas (CPRIT RP210045). Z.Z. was partially supported by NIH grants (R01LM012806, U01AG079847, and R01CA276513) and Cancer Prevention and Research Institute of Texas (CPRIT RP180734). We thank the authors of Immune Dictionary for providing the high-quality reference dataset that was used for training Scupa, and thank the authors of UCE for providing the foundation model that Scupa depends on.

## Author contributions

W.L. and Z.Z. conceived the project and wrote the manuscript. W.L. collected data, developed the method, and performed the analysis.

