## Supplementary Figures for "Scupa: Single-cell unified polarization assessment of immune cells using the single-cell foundation model"

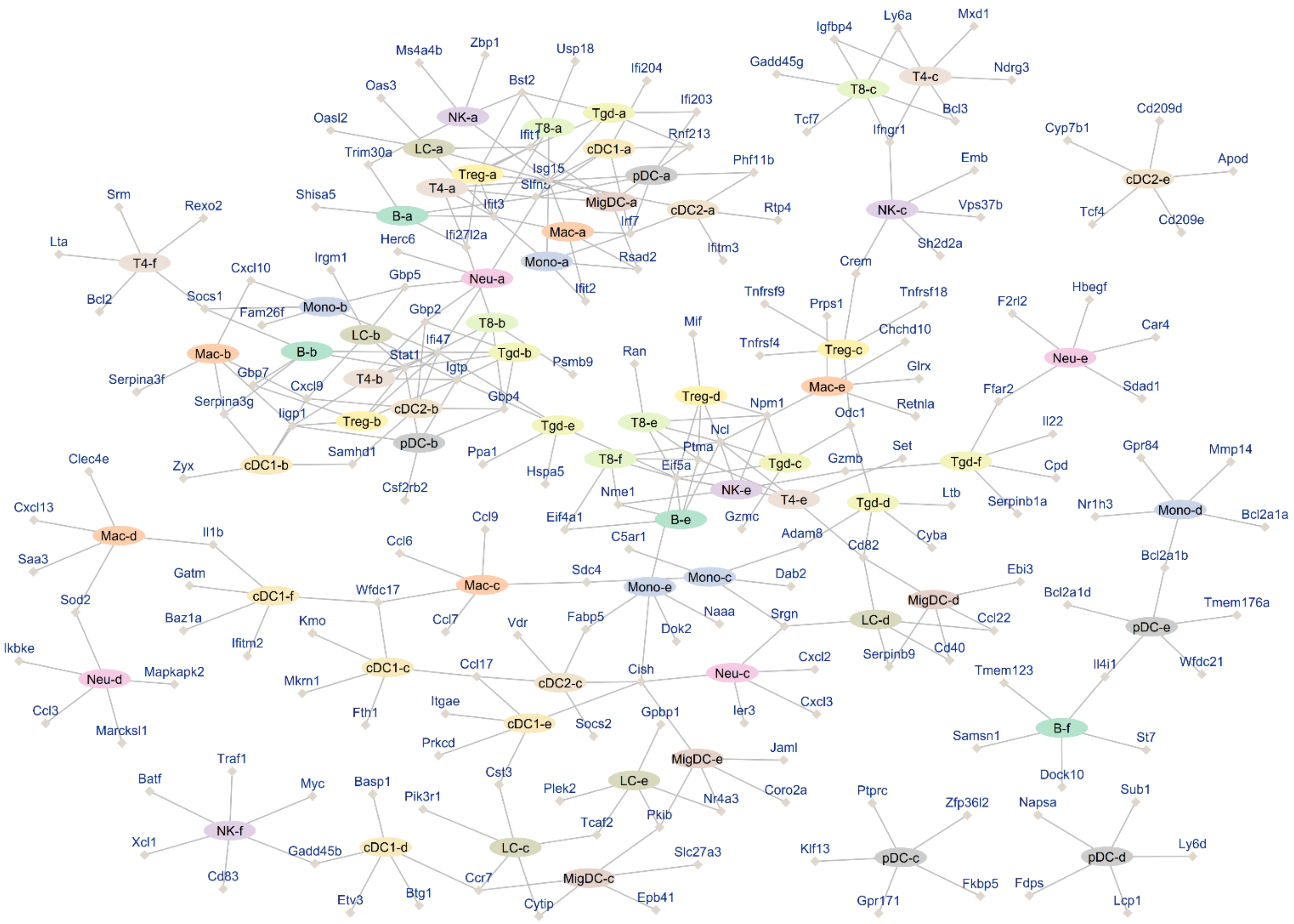


**Supplementary Fig. 1. A network showing the immune cell polarization states with top five marker genes.**

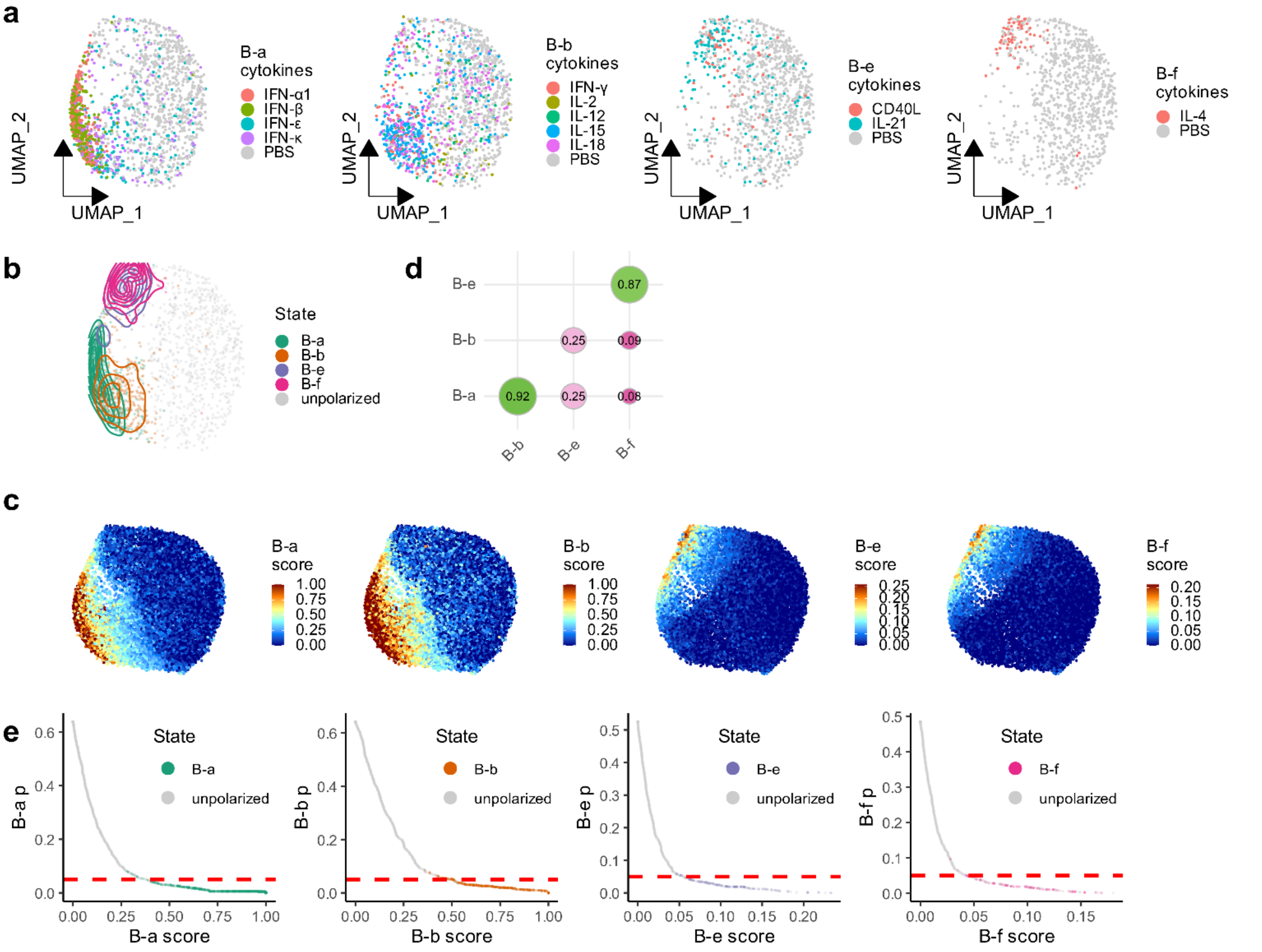


**Supplementary Fig. 2. Scupa learns the representation of polarization states in B cells.** a. UMAP plots showing B cells from the samples treated with driving cytokines of each polarization state or from the control samples treated with PBS. UMAPs are derived from UCE cell embeddings rather than gene expression. b. UMAP plot showing the distribution of fully polarized cells of each polarization state. c. UMAP plots showing polarization scores of each polarization state from Scupa prediction. d. The Spearman correlation coefficients between polarization scores from each two polarization states. e. The polarization scores and p-values of fully polarized cells of each polarization state and unpolarized cells. The red dashed line indicates p-value=0.05.

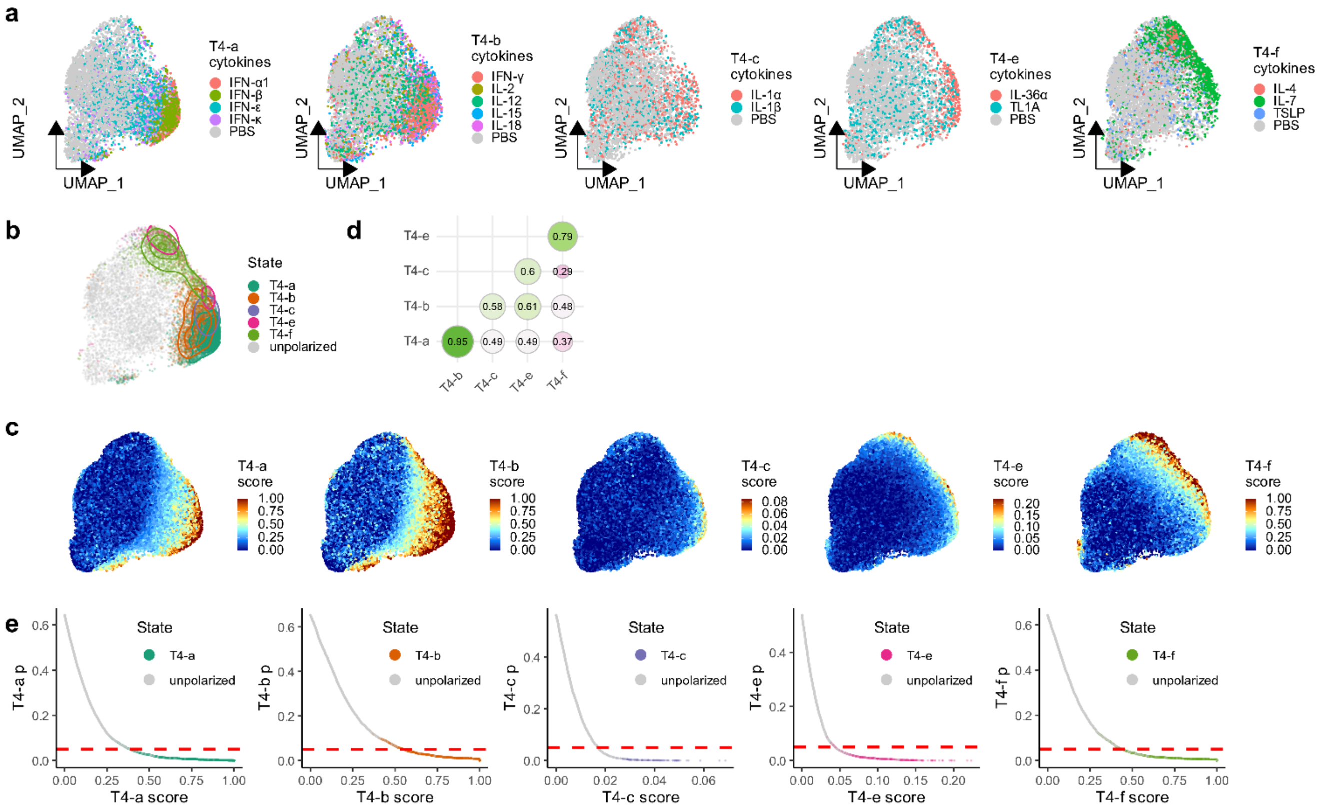


**Supplementary Fig. 3. Scupa learns the representation of polarization states in CD4$^+$ T cells.** a. UMAP plots showing CD4$^+$ T cells from the samples treated with driving cytokines of each polarization state or from the control samples treated with PBS. UMAPs are derived from UCE cell embeddings rather than gene expression. b. UMAP plot showing the distribution of fully polarized cells of each polarization state. c. UMAP plots showing polarization scores of each polarization state from Scupa prediction. d. The Spearman correlation coefficients between polarization scores from each two polarization states. e. The polarization scores and p-values of fully polarized cells of each polarization state and unpolarized cells. The red dashed line indicates p-value=0.05.

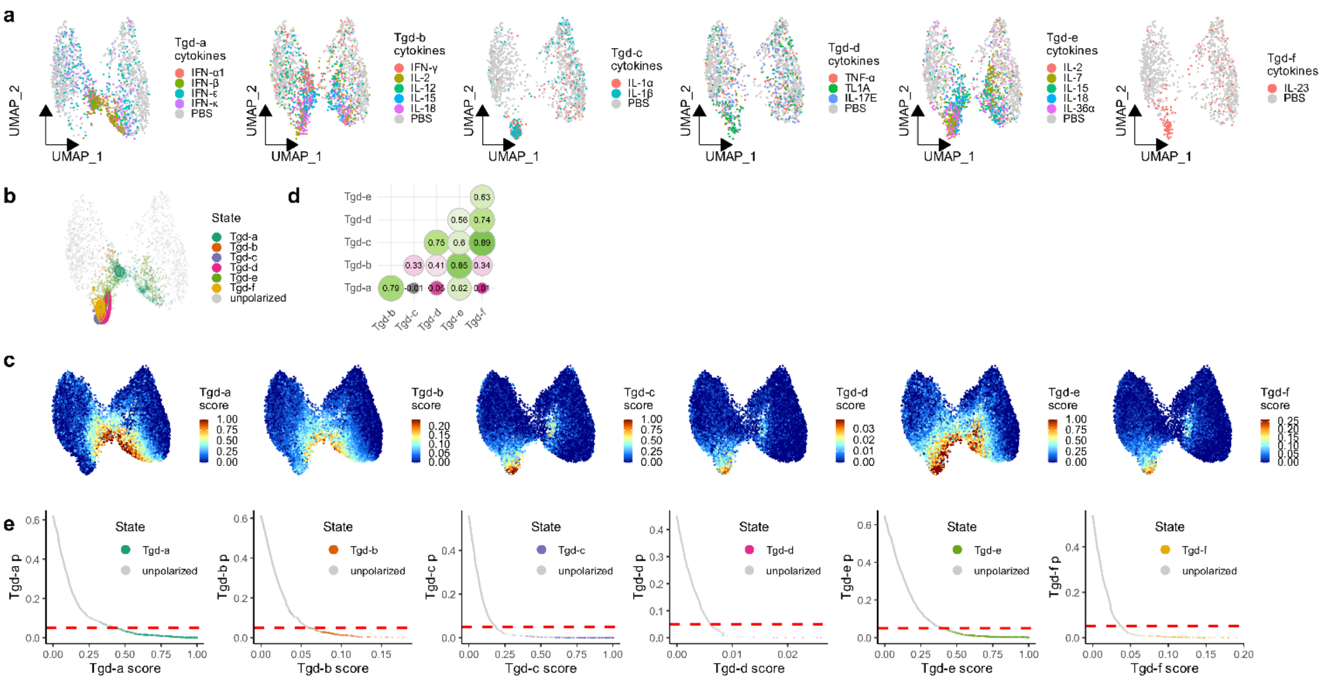

**Supplementary Fig. 4. Scupa learns the representation of polarization states in gamma-delta T (Tgd) cells.** a. UMAP plots showing Tgd cells from the samples treated with driving cytokines of each polarization state or from the control samples treated with PBS. UMAPs are derived from UCE cell embeddings rather than gene expression. b. UMAP plot showing the distribution of fully polarized cells of each polarization state. c. UMAP plots showing polarization scores of each polarization state from Scupa prediction. d. The Spearman correlation coefficients between polarization scores from each two polarization states. e. The polarization scores and p-values of fully polarized cells of each polarization state and unpolarized cells. The red dashed line indicates p-value=0.05.

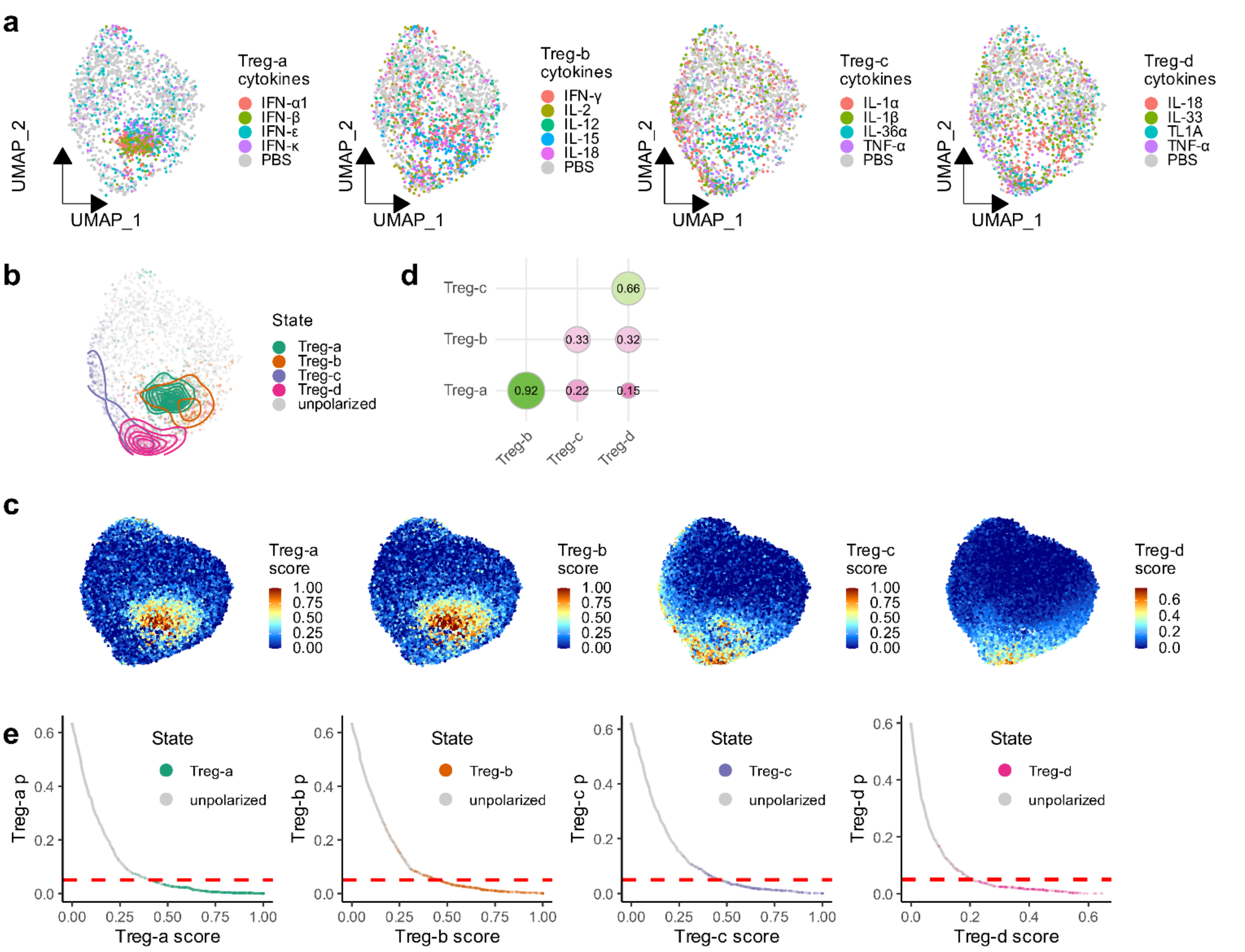


**Supplementary Fig. 5. Scupa learns the representation of polarization states in regulatory T (Treg) cells.** a. UMAP plots showing Treg cells from the samples treated with driving cytokines of each polarization state or form the control samples treated with PBS. UMAPs are derived from UCE cell embeddings rather than gene expression. b. UMAP plot showing the distribution of fully polarized cells of each polarization state. c. UMAP plots showing polarization scores of each polarization state from Scupa prediction. d. The Spearman correlation coefficients between polarization scores from each two polarization states. e. The polarization scores and p-values of fully polarized cells of each polarization state and unpolarized cells. The red dashed line indicates p-value=0.05.

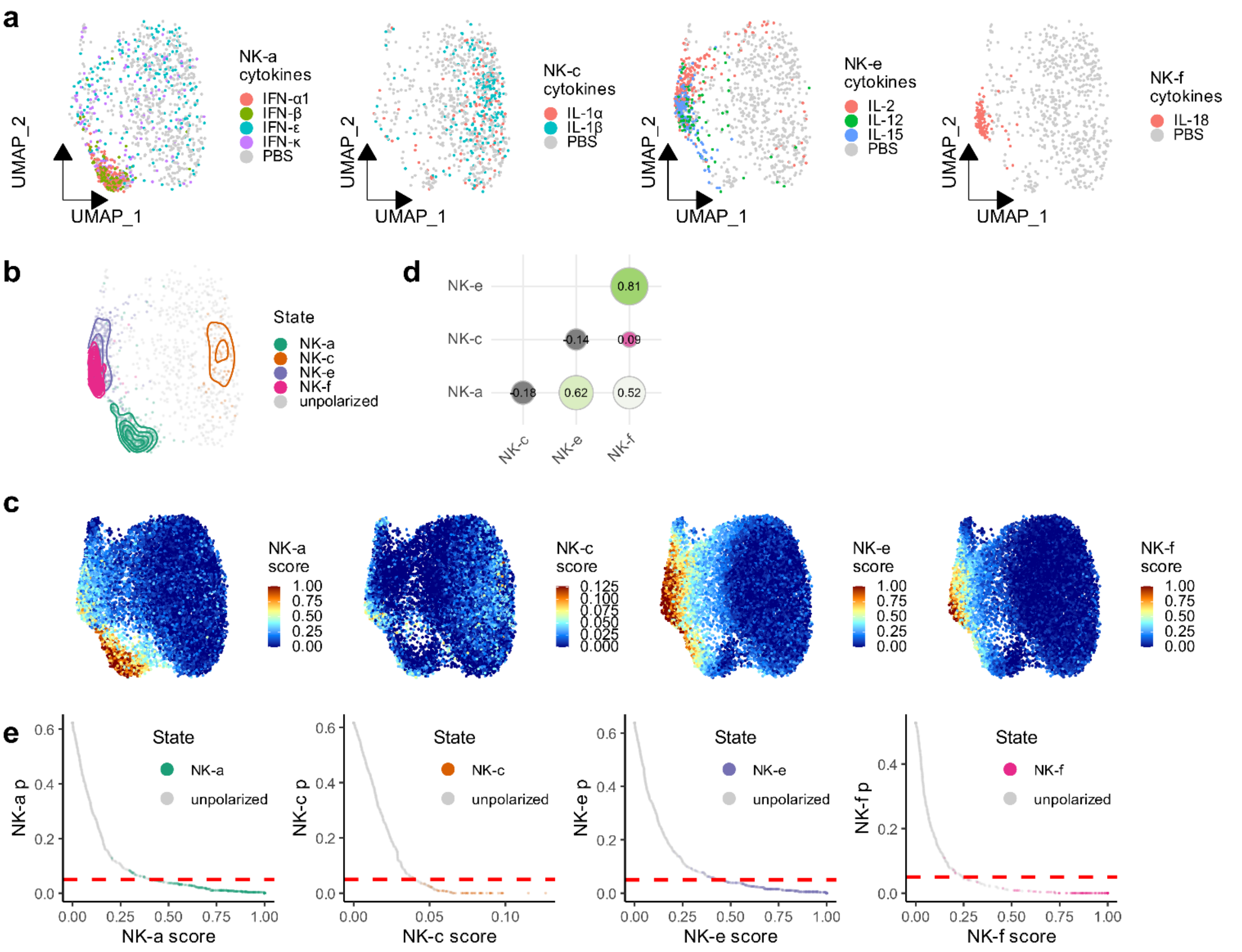


**Supplementary Fig. 6. Scupa learns the representation of polarization states in NK cells.** a. UMAP plots showing NK cells from the samples treated with driving cytokines of each polarization state or from the control samples treated with PBS. UMAPs are derived from UCE cell embeddings rather than gene expression. b. UMAP plot showing the distribution of fully polarized cells of each polarization state. c. UMAP plots showing polarization scores of each polarization state from Scupa prediction. d. The Spearman correlation coefficients between polarization scores from each two polarization states. e. The polarization scores and p-values of fully polarized cells of each polarization state and unpolarized cells. The red dashed line indicates p-value=0.05.

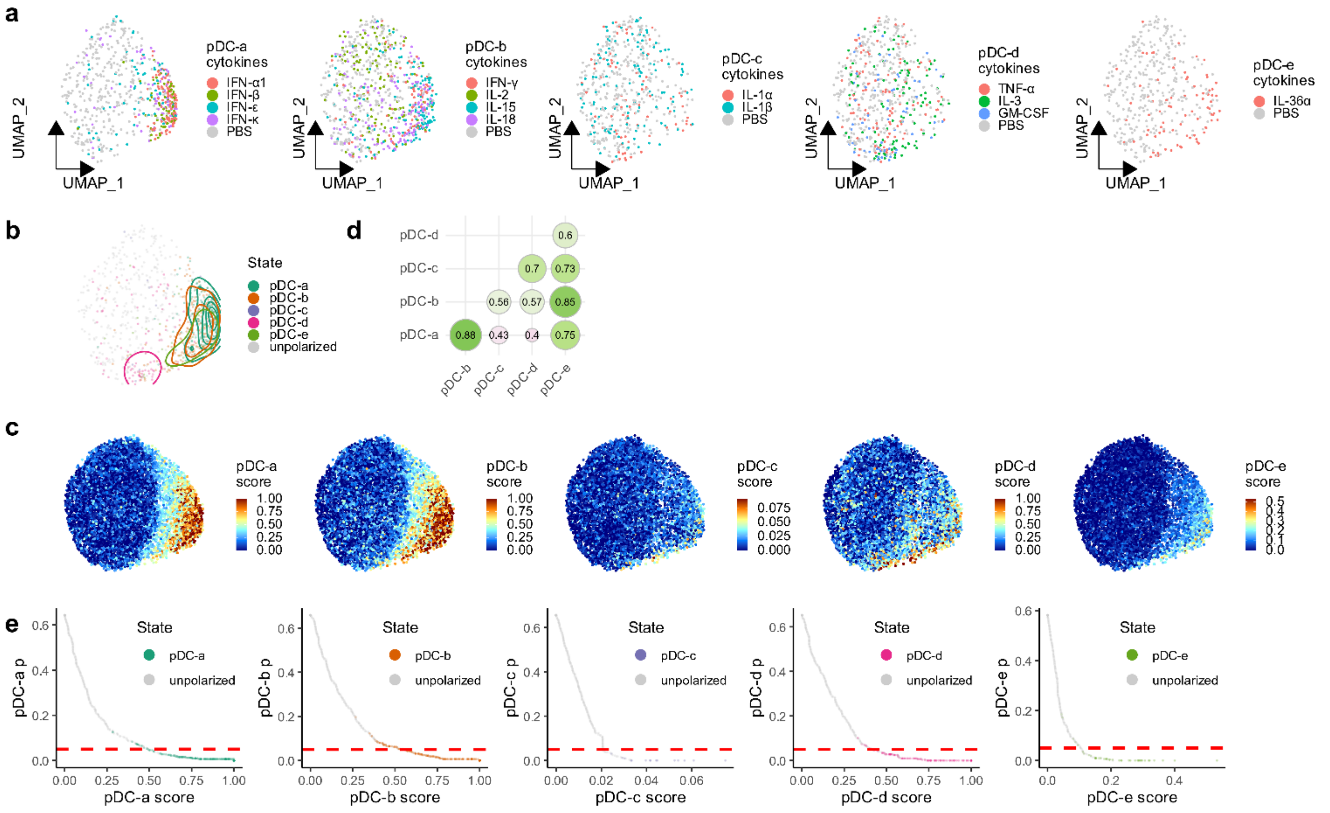


**Supplementary Fig. 7. Scupa learns the representation of polarization states in plasmacytoid dendritic cells (pDCs).** a. UMAP plots showing pDCs from the samples treated with driving cytokines of each polarization state or from the control samples treated with PBS. UMAPs are derived from UCE cell embeddings rather than gene expression. b. UMAP plot showing the distribution of fully polarized cells of each polarization state. c. UMAP plots showing polarization scores of each polarization state from Scupa prediction. d. The Spearman correlation coefficients between polarization scores from each two polarization states. e. The polarization scores and p-values of fully polarized cells of each polarization state and unpolarized cells. The red dashed line indicates p-value=0.05.

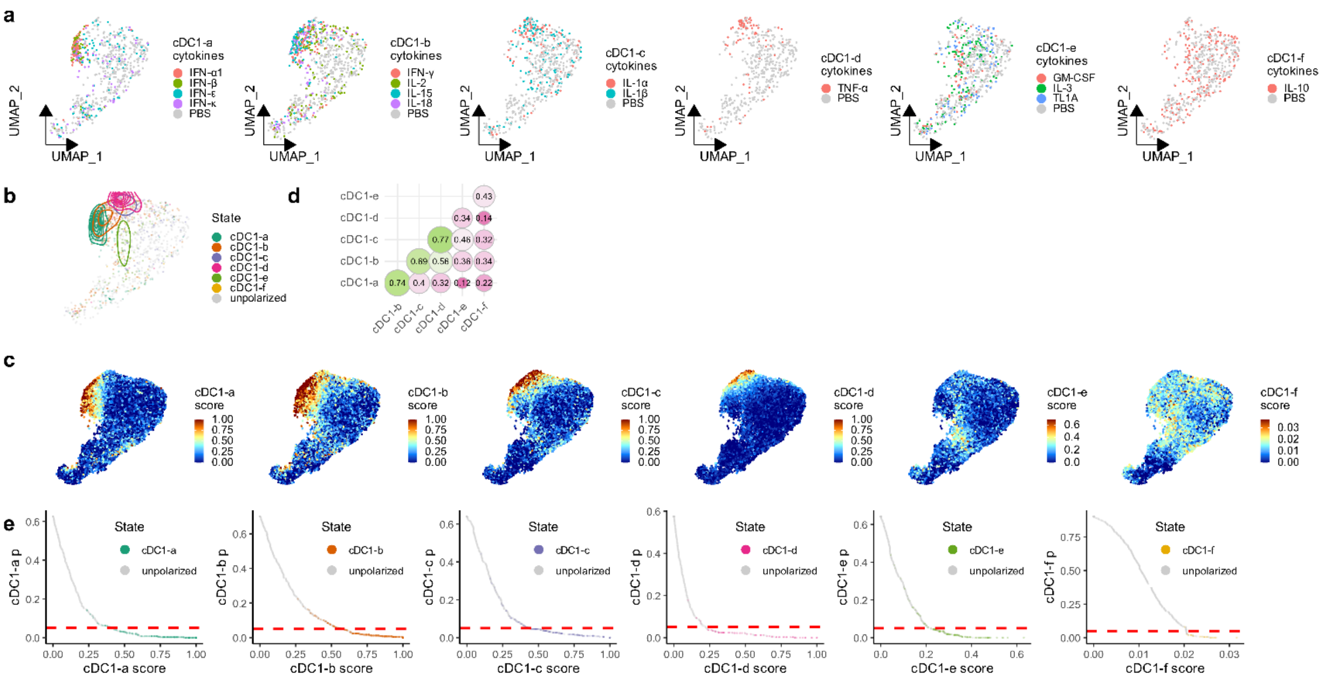


**Supplementary Fig. 8. Scupa learns the representation of polarization states in conventional dendritic cells 1 (cDC1).** a. UMAP plots showing cDC1 from the samples treated with driving cytokines of each polarization state or from the control samples treated with PBS. UMAPs are derived from UCE cell embeddings rather than gene expression. b. UMAP plot showing the distribution of fully polarized cells of each polarization state. c. UMAP plots showing polarization scores of each polarization state from Scupa prediction. d. The Spearman correlation coefficients between polarization scores from each two polarization states. e. The polarization scores and p-values of fully polarized cells of each polarization state and unpolarized cells. The red dashed line indicates p-value=0.05.

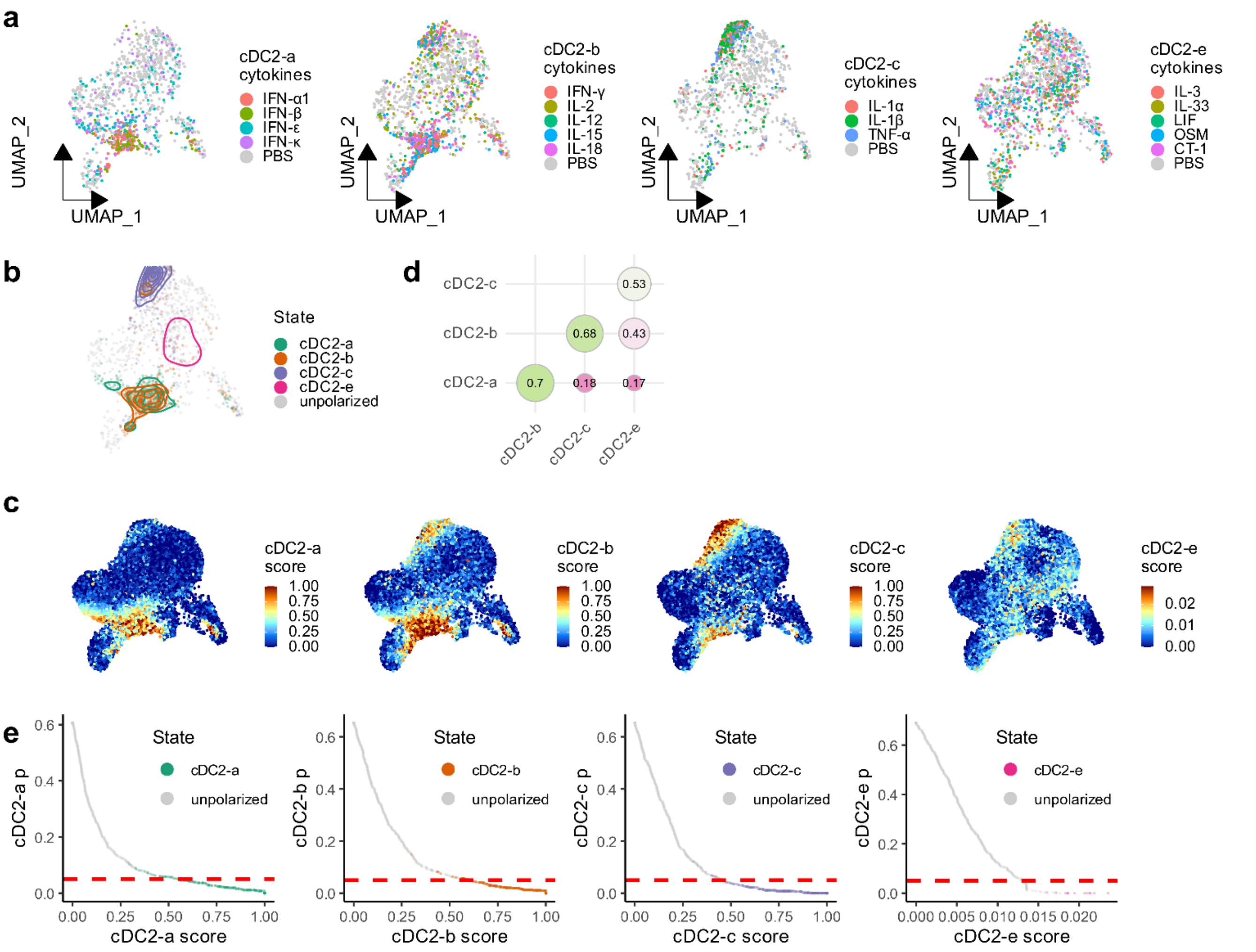


**Supplementary Fig. 9. Scupa learns the representation of polarization states in conventional dendritic cells 2 (cDC2).** a. UMAP plots showing cDC2 from the samples treated with driving cytokines of each polarization state or from the control samples treated with PBS. UMAPs are derived from UCE cell embeddings rather than gene expression. b. UMAP plot showing the distribution of fully polarized cells of each polarization state. c. UMAP plots showing polarization scores of each polarization state from Scupa prediction. d. The Spearman correlation coefficients between polarization scores from each two polarization states. e. The polarization scores and p-values of fully polarized cells of each polarization state and unpolarized cells. The red dashed line indicates p-value=0.05.

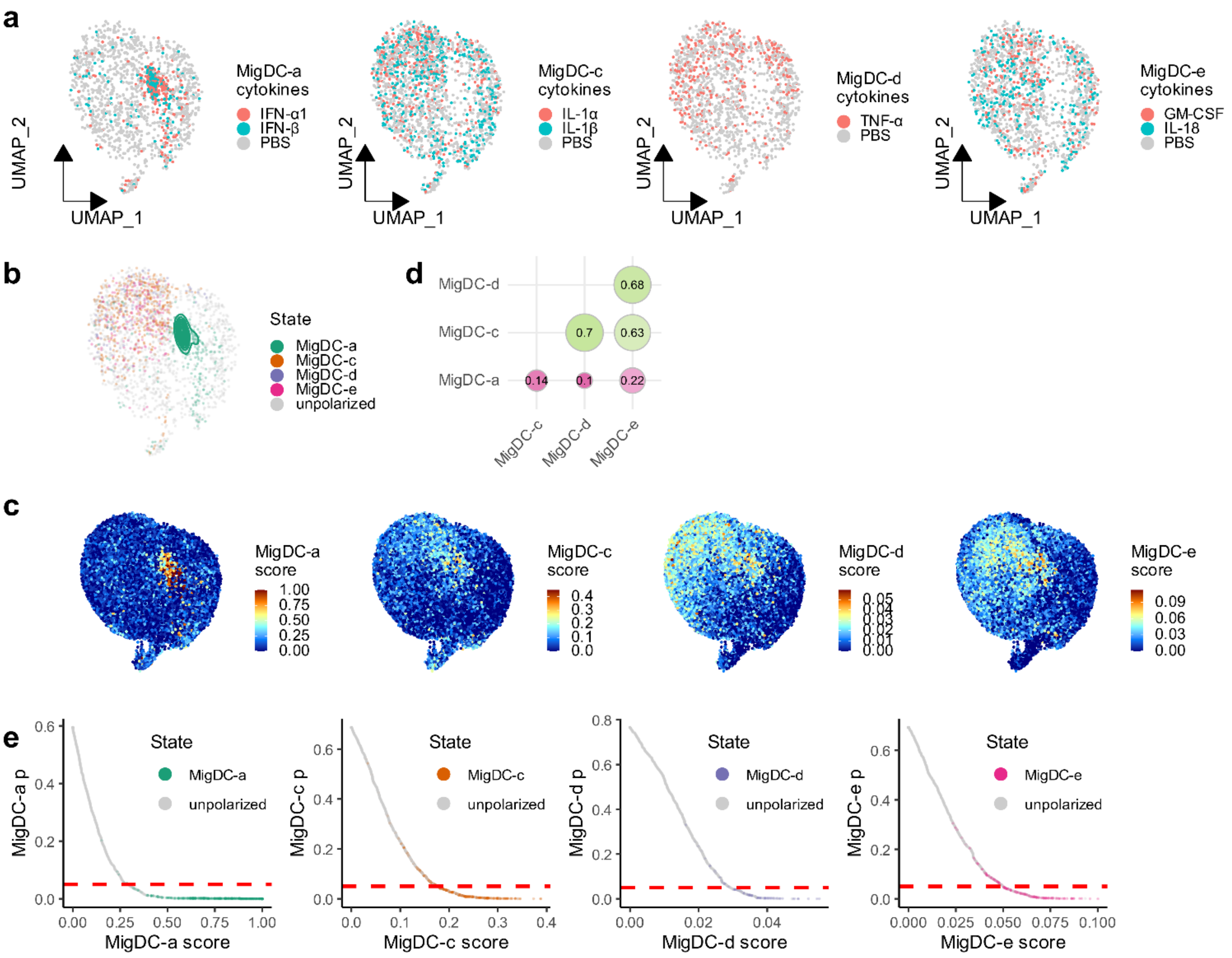


**Supplementary Fig. 10. Scupa learns the representation of polarization states in migratory dendritic cells (MigDCs).** a. UMAP plots showing MigDCs from the samples treated with driving cytokines of each polarization state or from the control samples treated with PBS. UMAPs are derived from UCE cell embeddings rather than gene expression. b. UMAP plot showing the distribution of fully polarized cells of each polarization state. c. UMAP plots showing polarization scores of each polarization state from Scupa prediction. d. The Spearman correlation coefficients between polarization scores from each two polarization states. e. The polarization scores and p-values of fully polarized cells of each polarization state and unpolarized cells. The red dashed line indicates p-value=0.05.

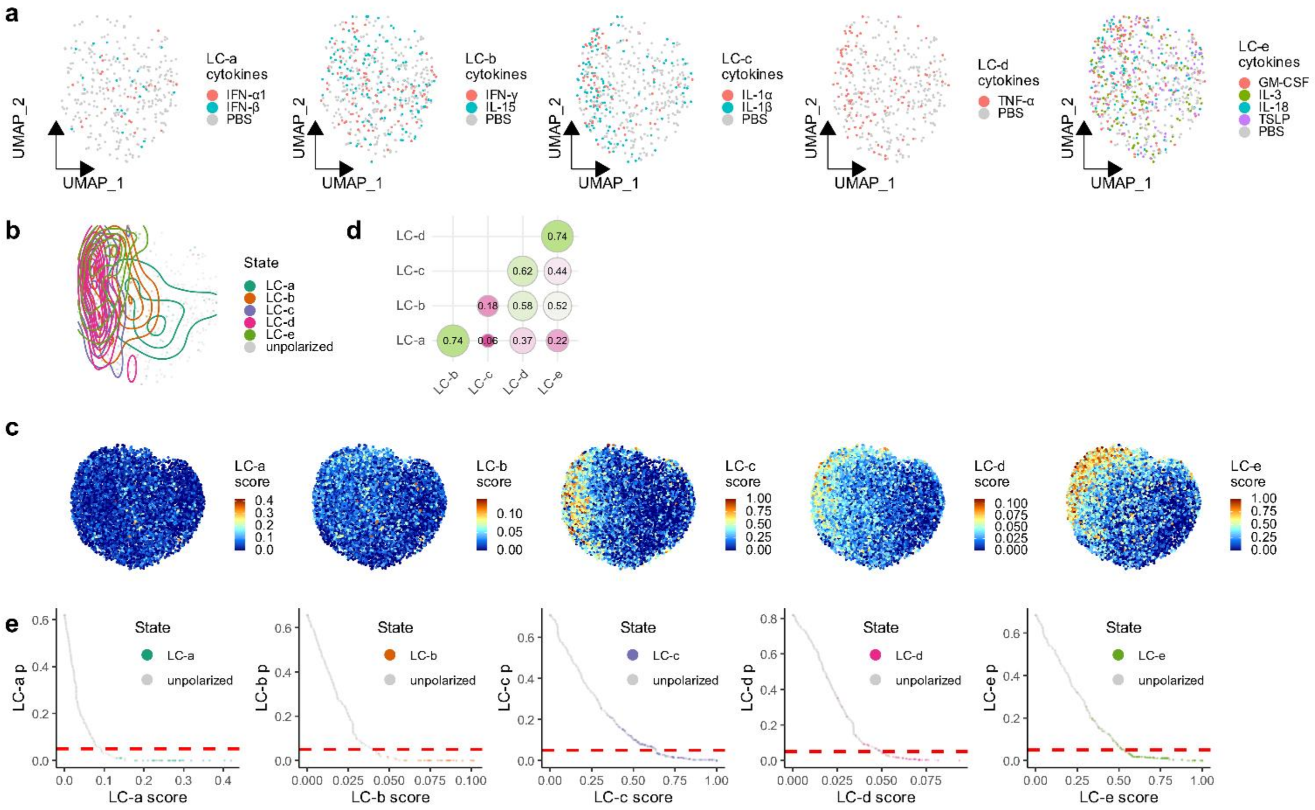

**Supplementary Fig. 11. Scupa learns the representation of polarization states in Langerhans cells (LCs).** a. UMAP plots showing LCs from the samples treated with driving cytokines of each polarization state or from the control samples treated with PBS. UMAPs are derived from UCE cell embeddings rather than gene expression. b. UMAP plot showing the distribution of fully polarized cells of each polarization state. c. UMAP plots showing polarization scores of each polarization state from Scupa prediction. d. The Spearman correlation coefficients between polarization scores from each two polarization states. e. The polarization scores and p-values of fully polarized cells of each polarization state and unpolarized cells. The red dashed line indicates p-value=0.05.

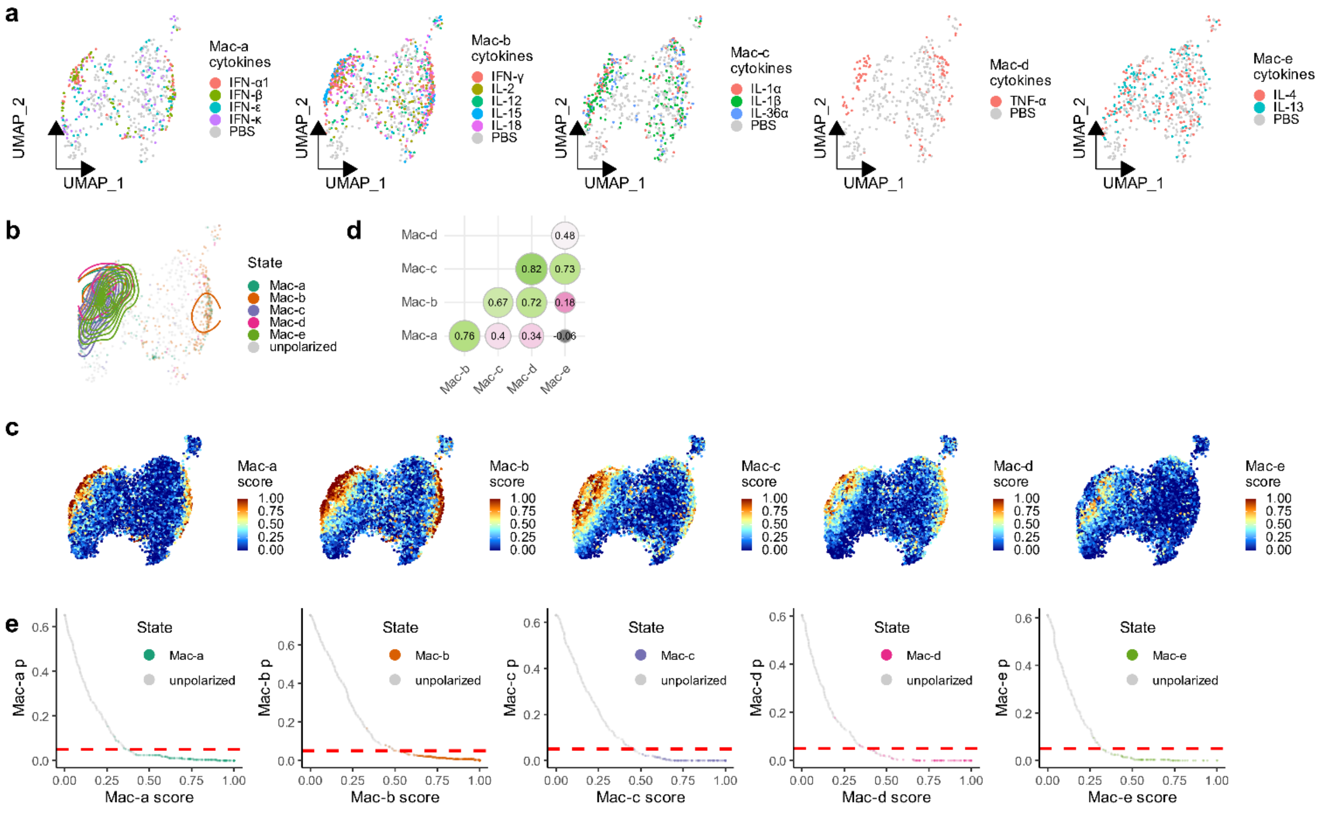


**Supplementary Fig. 12. Scupa learns the representation of polarization states in macrophages.** a. UMAP plots showing macrophages from the samples treated with driving cytokines of each polarization state or from the control samples treated with PBS. UMAPs are derived from UCE cell embeddings rather than gene expression. b. UMAP plot showing the distribution of fully polarized cells of each polarization state. c. UMAP plots showing polarization scores of each polarization state from Scupa prediction. d. The Spearman correlation coefficients between polarization scores from each two polarization states. e. The polarization scores and p-values of fully polarized cells of each polarization state and unpolarized cells. The red dashed line indicates p-value=0.05.

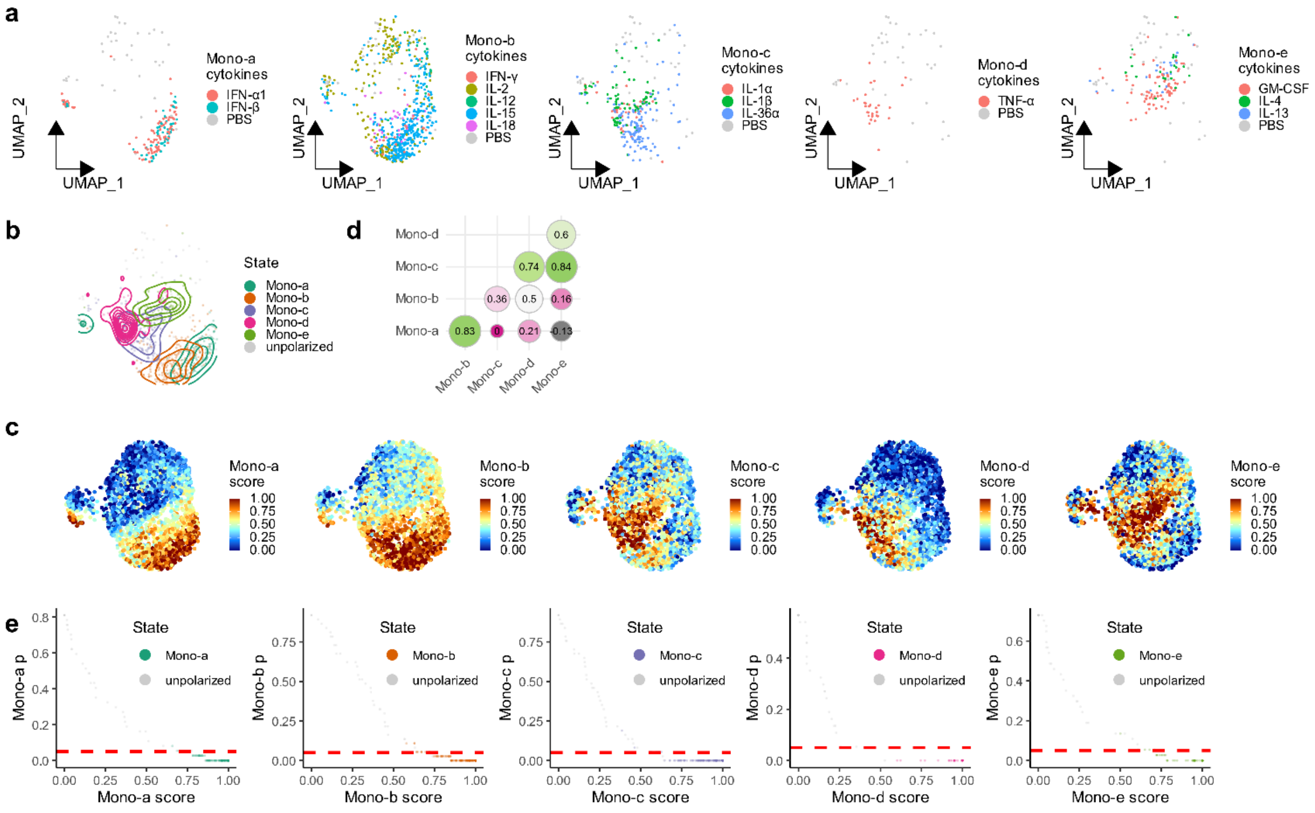


**Supplementary Fig. 13. Scupa learns the representation of polarization states in monocytes.** a. UMAP plots showing monocytes from the samples treated with driving cytokines of each polarization state or from the control samples treated with PBS. UMAPs are derived from UCE cell embeddings rather than gene expression. b. UMAP plot showing the distribution of fully polarized cells of each polarization state. c. UMAP plots showing polarization scores of each polarization state from Scupa prediction. d. The Spearman correlation coefficients between polarization scores from each two polarization states. e. The polarization scores and p-values of fully polarized cells of each polarization state and unpolarized cells. The red dashed line indicates p-value=0.05.

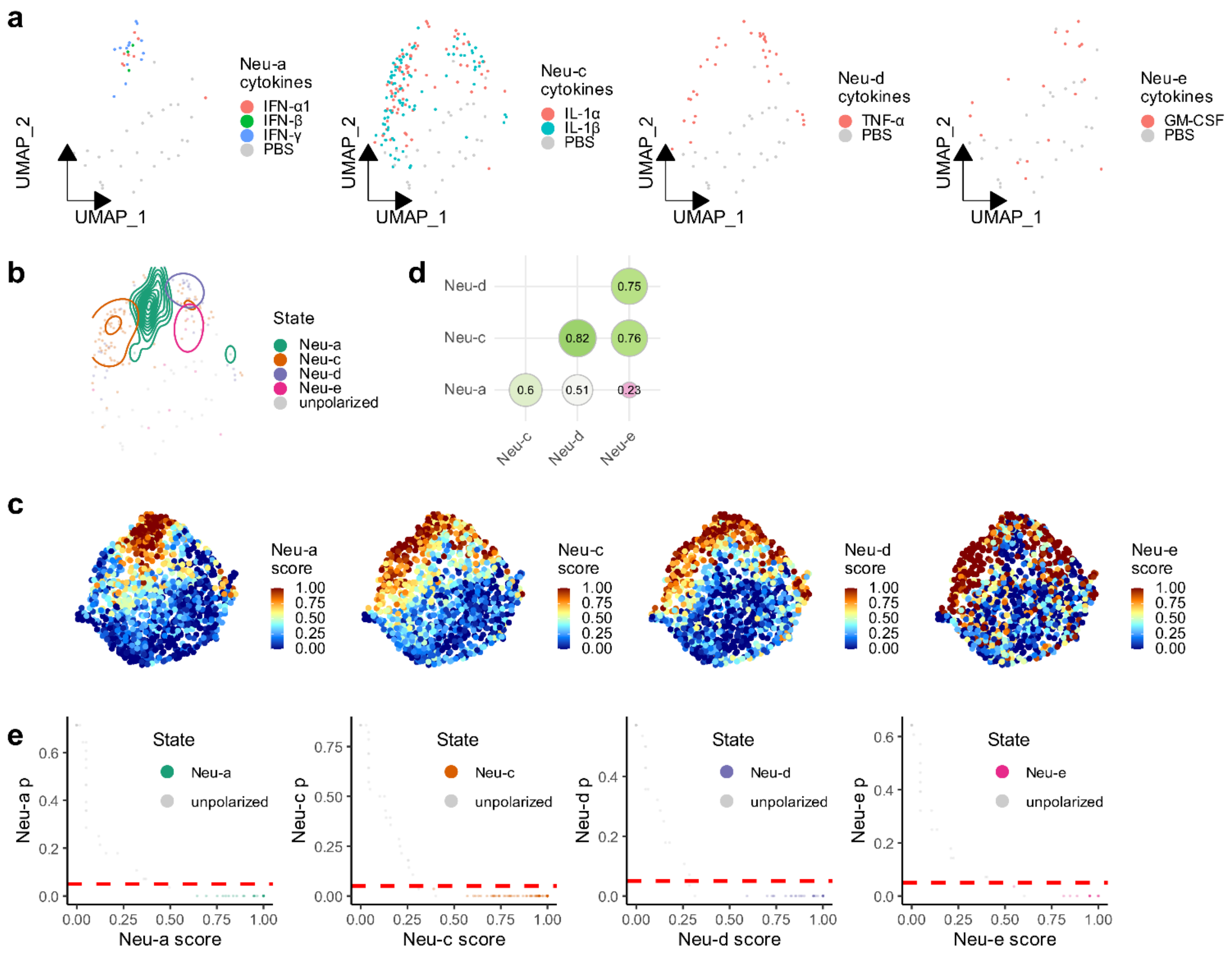


**Supplementary Fig. 14. Scupa learns the representation of polarization states in neutrophils.** a. UMAP plots showing neutrophils from the samples treated with driving cytokines of each polarization state or from the control samples treated with PBS. UMAPs are derived from UCE cell embeddings rather than gene expression. b. UMAP plot showing the distribution of fully polarized cells of each polarization state. c. UMAP plots showing polarization scores of each polarization state from Scupa prediction. d. The Spearman correlation coefficients between polarization scores from each two polarization states. e. The polarization scores and p-values of fully polarized cells of each polarization state and unpolarized cells. The red dashed line indicates p-value=0.05.

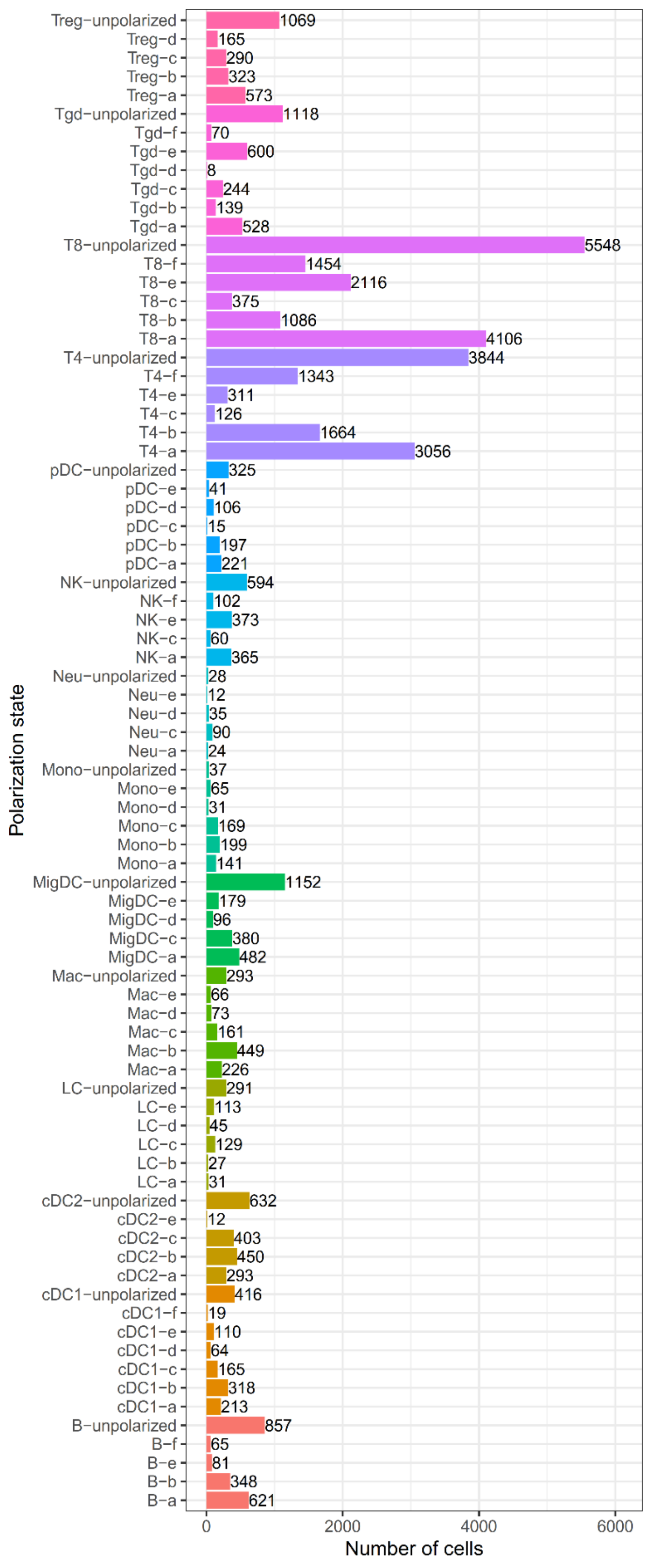


**Supplementary Fig. 15. The number of cells in each immune cell polarization state from the Immune Dictionary.**

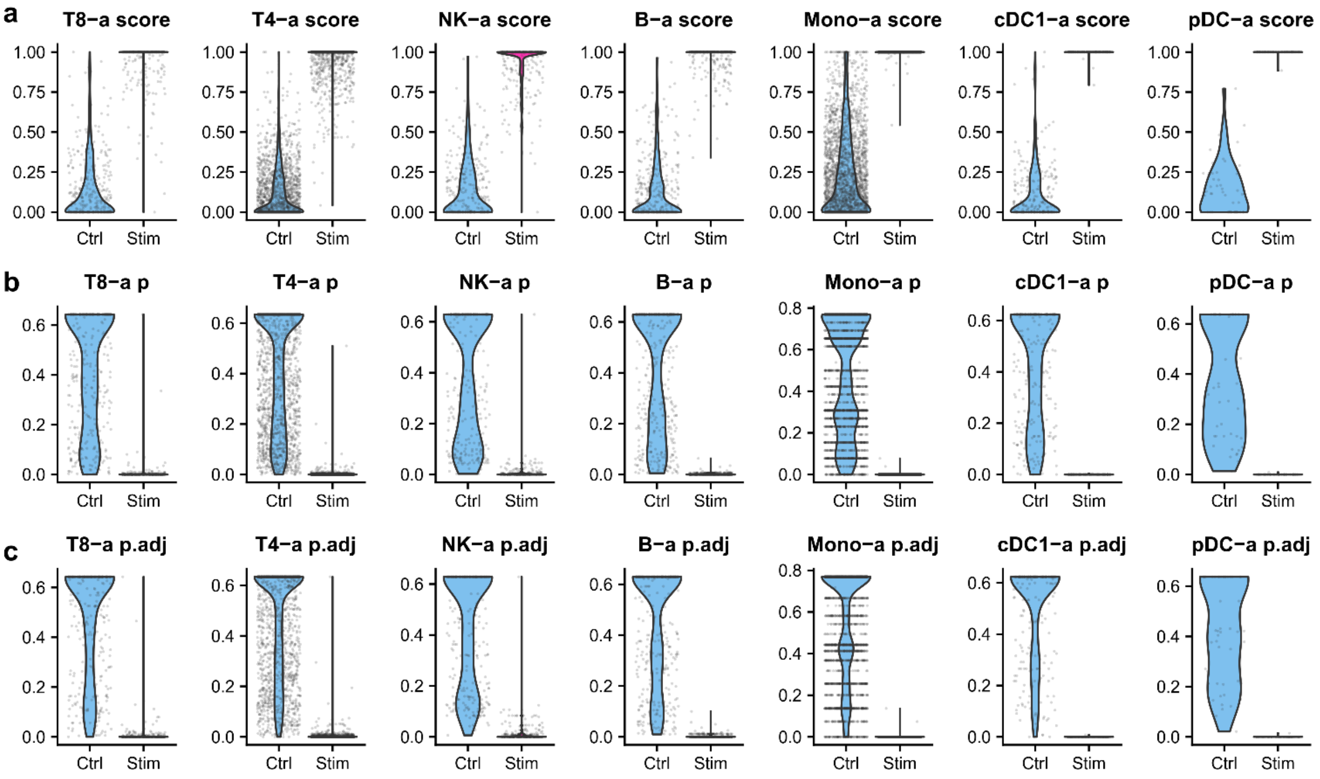


**Supplementary Fig. 16. Scupa classified polarized cells and unpolarized cells in IFN-β treatment.** a-c. Immune cells from the control and IFN-β-stimulated samples showed significant differences in polarization score distribution (a), p-value distribution (b), and adjusted p-value distribution (c). Ctrl: untreated control sample. Stim: interferon-stimulated sample.
